# DOT1L regulates chromatin reorganization and gene expression during sperm differentiation

**DOI:** 10.1101/2022.10.17.512530

**Authors:** Mélina Blanco, Laila El Khattabi, Clara Gobé, Marion Crespo, Manon Coulée, Alberto de la Iglesia, Côme Ialy-Radio, Clementine Lapoujade, Maëlle Givelet, Marion Delessard, Ivan Seller-Corona, Kosuke Yamaguchi, Nadège Vernet, Fred Van Leeuwen, Alban Lermine, Yuki Okada, Romain Daveau, Rafael Oliva, Pierre Fouchet, Ahmed Ziyyat, Delphine Pflieger, Julie Cocquet

**Author notes:** equal contribution (Laila El Khattabi, Clara Gobé).

## Abstract

Spermatozoa have a unique genome organization: their chromatin is almost completely devoid of histones and is formed instead of protamines which confer a high level of compaction and preserve paternal genome integrity until fertilization. Histone-to-protamine transition takes place in spermatids and is indispensable for the production of functional sperm. Here we show that the H3K79-methyltransferase DOT1L controls spermatid chromatin remodelling and subsequent reorganization and compaction of spermatozoon genome. Using a mouse model in which *Dot1l* is knocked-out (KO) in postnatal male germ cells, we found that *Dot1l*-KO sperm chromatin is less compact and has an abnormal content, characterized by the presence of transition proteins, immature protamine 2 forms and a higher level of histones. Proteomics and transcriptomics analyses performed on spermatids reveal that *Dot1l*-KO modifies the chromatin prior to histone removal, and leads to the deregulation of genes involved in flagellum formation and apoptosis during spermatid differentiation. As a consequence of these chromatin and gene expression defects, *Dot1l*-KO spermatozoa have less compact heads and are less motile which results in impaired fertility.

## Introduction

During the postmeiotic phase of spermatogenesis, male germ cells known as spermatids undergo profound morphological and functional changes to differentiate in spermatozoa. This process is driven by a rich genetic program characterized by the expression of thousands of genes in round spermatids (Chen *et al*, 2018; da Cruz *et al*, 2016; Ernst *et al*, 2019; Green *et al*, 2018; Soumillon *et al*, 2013). Then, when spermatids elongate, their chromatin is extensively remodeled which ultimately results in the eviction of ∼85-99 % of nuclear histones and their replacement by more basic, smaller proteins, called protamines [for reviews, see (Bao & Bedford, 2016; Rathke *et al*, 2014)]. This unique chromatin reorganization induces a 6-10 times higher level of compaction of the spermatozoon chromatin compared to the canonical nucleosome-based chromatin (Ward & Coffey, 1991). Non-histone packaging of male germ cell genome is conserved throughout evolution and expected to be important to protect paternal DNA from damages and prepare its reprograming in the zygote, if fertilization occurs (Rathke *et al*., 2014). A compact genome is also an advantage for the motile spermatozoa as it is compatible with a smaller and more hydrodynamic head (Braun, 2001).

In the past decades, the molecular mechanisms driving histone eviction and subsequent compaction of the sperm genome have been the focus of several studies which have demonstrated the essential role of transition proteins (Zhao *et al*, 2004a; Zhao *et al*, 2004b), of histone variants (Barral *et al*, 2017; Montellier *et al*, 2013), of histone post-translational modifications (PTMs), in particular hyperacetylation (Goudarzi *et al*, 2016; Oliva *et al*, 1987), and of writers and readers of histone acetylation (Dong *et al*, 2017; Gaucher *et al*, 2012; Luense *et al*, 2019; Shang *et al*, 2007; Shiota *et al*, 2018). Interestingly, high levels of histone H3 Lysine 79 di- and tri-methylation (H3K79me2 and me3) have been observed at the same time as histone hyperacetylation, just prior to histone removal (Dottermusch-Heidel *et al*, 2014a; Dottermusch-Heidel *et al*, 2014b; Moretti *et al*, 2017) (Fig 1A), but their biological significance remains to date unknown. H3K79 methylation is mediated by one enzyme, encoded by the gene *Dot1l*, of which pattern of expression and sequence are conserved from Drosophila to mammals. *Dot1l* has been implicated in development, cell reprogramming, differentiation and proliferation [for review, see (Kim *et al*, 2014; Vlaming & van Leeuwen, 2016)]. It has recently been found to be essential for spermatogonial stem cell self-renewal (Lin *et al*, 2022) but its role in postmeiotic male germ cells, where it is the most highly expressed (Dottermusch-Heidel *et al*., 2014a; Dottermusch-Heidel *et al*., 2014b; Moretti *et al*., 2017), has not been addressed.

**Figure 1.**
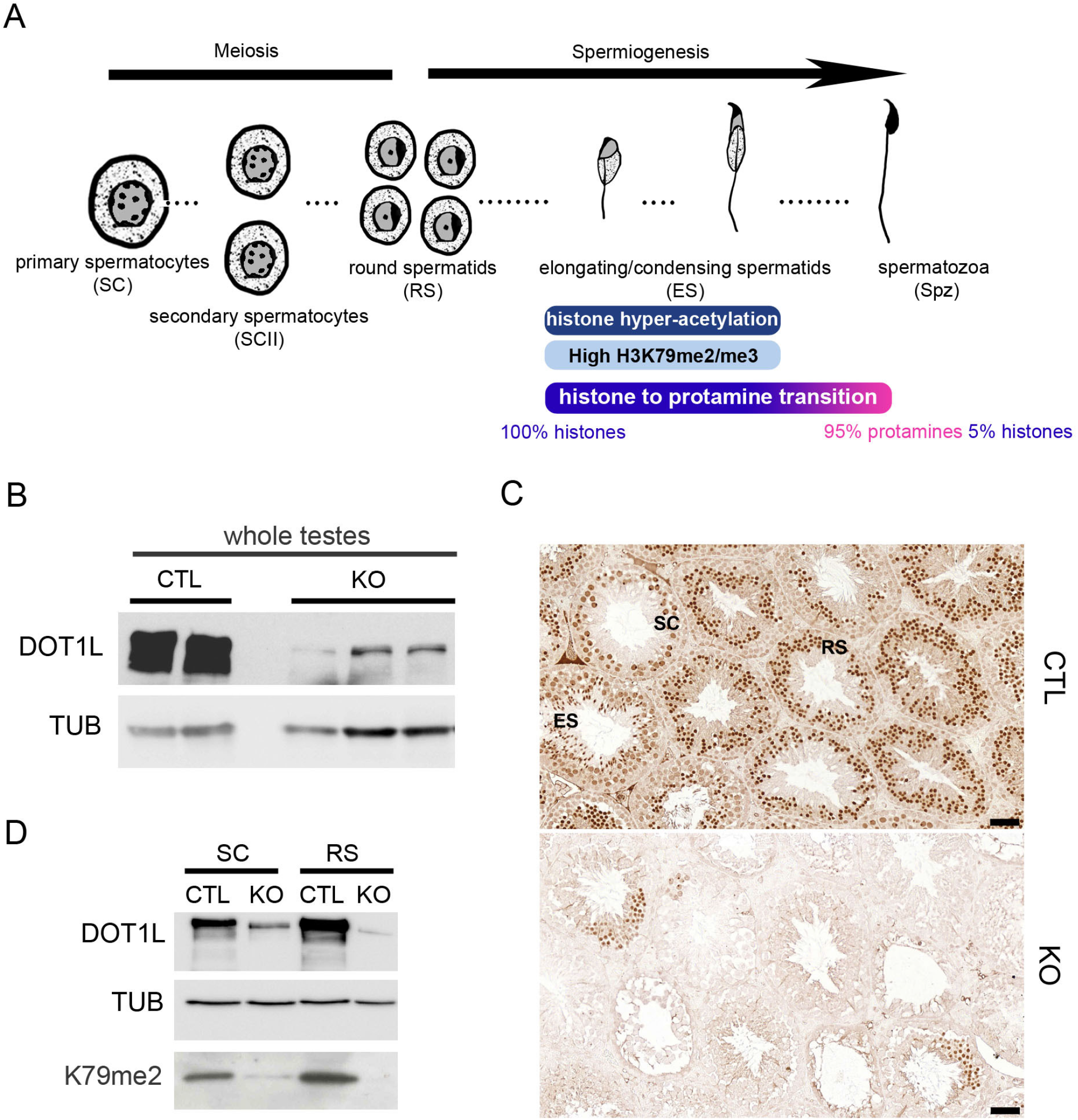
DOT1L expression in meiotic and postmeiotic germ cells of *Dot1l*-KO and CTL males. A) Spermatogenesis scheme representing the dynamic of histone to protamine transition. B) Western blot detection of DOT1L and TUBULIN (TUB) in whole testicular protein extracts from CTL and *Dot1l*-KO (KO) adult mice. C) Immunohistochemistry detection of DOT1L in testicular sections from CTL and *Dot1l*-KO (KO) adult mice. SC = Spermatocytes, RS = Round spermatids, ES = Elongating spermatids. Pictures were taken using the same parameters. Scale bars indicate 50µm. D) Western blot detection of DOT1L and H3K79me2 in *Dot1l*-KO (KO) and CTL germ cells. Normalization was performed by detecting the membrane with anti-TUBULIN (TUB). SC = primary spermatocytes, RS = round spermatids. See also Fig EV1 and Appendix FigS1.

In the present paper, we investigated *Dot1l* role in postmeiotic cells by a comprehensive analysis of the phenotypic and molecular consequences of its knockout in mouse male germ cells. We found that *Dot1l* is required for gene regulation and chromatin reorganization in spermatids and that its knockout leads to the production of abnormally shaped nonfunctional spermatozoa with a deficient chromatin and nucleus compaction.

## Results

### *Dot1l* is essential for spermatogenesis and male fertility

In both mouse and human, *Dot1l/DOT1L* gene is highly expressed in the testis (Appendix FigS1A). More precisely, it is expressed in all testicular cells with a strong expression at postnatal stage in meiotic cells (spermatocytes) and postmeiotic cells (spermatids) (Appendix FigS1A-C) (Dottermusch-Heidel *et al*., 2014a; Dottermusch-Heidel *et al*., 2014b; Green *et al*., 2018; Moretti *et al*., 2017). To investigate the role of *Dot1l* in male germ cells, we generated a conditional knockout using *Stra8-Cre r*ecombinase (Sadate-Ngatchou *et al*, 2008) and a mouse transgenic line in which *Dot1l* exon 2 is flanked by LoxP sites (Jo *et al*, 2011) (Appendix FigS1D). Previous studies have reported that *Stra8-Cre* is specifically expressed in the male germline where it is expressed from postnatal day 3 and CRE recombinase reaches maximum efficiency in pachytene spermatocytes (Bao *et al*, 2013; Sadate-Ngatchou *et al*., 2008). Bao *et al*. have also observed that *Stra8-Cre* is more efficient at cutting one allele rather than two (Bao *et al*., 2013). We therefore generated mice in which one allele of *Dot1l* is floxed and one allele is already deleted (Δ) to increase efficiency of floxed exon excision upon *Cre* recombinase expression (*Dot1l*^*Fl/Δ*^; *Stra8-Cre* mice, hereafter called *Dot1l*-KO or KO). First, to estimate the efficiency of *Dot1l* knockout, we detected DOTL1 protein by western blot and immunochemistry on adult testes from *Dot1l*-KO (*Dot1l*^*Fl/Δ*^; *Stra8-Cre*) and controls (*Dot1l*^*Fl/Fl*^) (Figs 1B-C and Appendix FigS1E-G). We found a reduction of >85 % of DOT1L signal but not complete abolishment/knockout of DOT1L level, despite strong activity of *Stra8-Cre* in most adult germ cells (Appendix FigS1H). Analyses of purified germ cell fractions confirmed reduced level but not complete absence of DOTL1 protein, along with a reduction of H3K79me2 level (Figs 1D, EV1 and Appendix FigS1I). Multiple protein isoforms of DOT1L have been described in the literature and several of them were observed in our western blots, including the canonical (∼165 kDa) and testis-specific (∼185 kDa) isoforms (Dottermusch-Heidel *et al*., 2014b; Zhang *et al*, 2004), as well as less studied isoforms such as Q679P5 (∼122KDa) or Q6XZL7 (∼68KDa) (https://www.uniprot.org). All of them were markedly reduced in *Dot1l*-KO testicular extracts (Appendix FigS1E and I).

To investigate the impact of *Dot1l* knockout on spermatogenesis and male fertility, the reproductive parameters of adult *Dot1l*-KO males were investigated: significant decreases in testis weight and sperm count were observed (independently of body weight, Fig 2A and Appendix FigS2A-B). Histological analyses revealed the presence of “empty” tubules without germ cells or missing germ cell layers (Fig 2B and Appendix FigS2C) highlighting a spermatogonial defect as described by Lin et al (Lin *et al*., 2022). However, there was no homogenous blockage/arrest at a specific stage of *Dot1l*-KO spermatogenesis.

**Figure 2.**
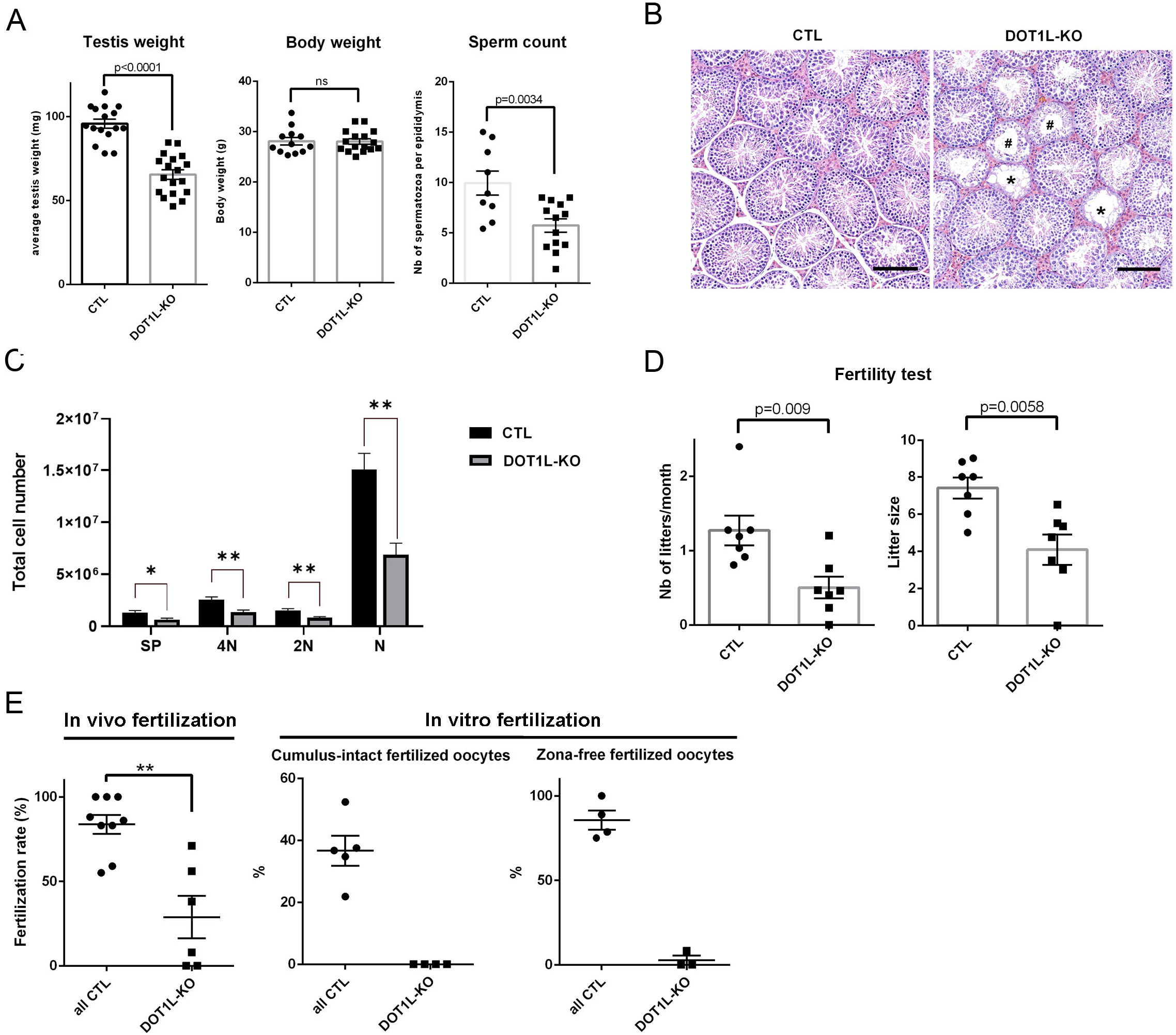
*Dot1l*-KO in male germ cells impairs spermatogenesis and male fertility. A) Scatter plots (mean + standard deviation) showing the average testis weight, body weight and sperm count in *Dot1l*-KO and CTL adult mice (2 to 4 month-old N>10 for CTL and *Dot1l*-KO samples). B) Histology of *Dot1l*-KO and CTL testes (periodic acid-Schiff staining). Scale bars indicate 200µm. C) Schematic diagram representing total number of each cell type per testis in CTL and *Dot1l*-KO males, as calculated by FACS. 4N = Primary spermatocytes, 2N = Secondary spermatocytes, N = spermatids, SP = Side Population representing premeiotic germ cells (spermatogonia). 8 CTL and 6 *Dot1l*-KO testes were analyzed. One or two stars indicate a p-value <0.05 or <0.01, respectively (Student t-test). D) Results from the tests of fertility (natural mating) of *Dot1l*-KO and CTL males (7 males in each group mated for ∼3 months with wild-type females). Schematic diagrams representing the number of litters and the litter size from progeny of *Dot1l*-KO or CTL males (mean + standard deviation). E) In vivo (left panel) and in vitro fertilization results. The percentages of fertilized oocytes are indicated for WT, CTL and HET males (‘all CTL’) and for *Dot1l*-KO males. The left panel shows the mean fertilization rate (%) of oocytes collected in superovulated females 24h after natural mating). Nine and 6 females with a positive vaginal plug were analyzed after mating with CTL and *Dot1l-*KO males respectively. This represents 113 and 73 oocytes analyzed after mating with CTL or KO males respectively. Two stars indicate a p-value <0.005 (unpaired t-test performed after angular transformation of percentages). The central and right panels show in vitro fertilization results using oocytes with intact cumulus, or using oocytes of which the zona pellucida was removed beforehand. See also Appendix FigS2.

These observations were confirmed by cytometry analyses in which the number of all germ cell populations (i.e. spermatogonia with Side Population phenotype, spermatocytes and spermatids) was found decreased in *Dot1l*-KO testes (Fig 2C), but their frequency was not altered (Appendix FigS2F). Collectively, these data show that the *Dot1l*-KO we produced does not lead to a spermatogenesis arrest. When mated to WT females, *Dot1l*-KO males were found to be hypofertile with significant reductions in the number of litters and in litter size compared to that of control (CTL) males (Fig 2D and Appendix FigS2D-E). We genotyped the few pups obtained and observed that the floxed allele was transmitted to 35% of babies (12 out of 34) and the deleted allele to 65% of them. This indicates that the deleted allele can be transmitted but less efficiently than what was expected based on our quantification of *Dot1l*-KO spermatids by immunofluorescence (i.e. only 65% of transmission of the deleted allele although >85% of spermatids were devoid of DOT1L protein).

The reduction in litter number and size could result from impaired fertilizing ability of *Dot1l*-KO sperm or from a developmental defect. To address this question, we measured the fertilization rate 24h after WT females were mated with *Dot1l*-KO or CTL males. The numbers of fertilized and non-fertilized oocytes collected in the oviduct ampulla were counted in both groups (a total of 113 oocytes were analyzed from 9 WT females mated with 4 CTL, and 73 oocytes from 6 WT females mated with 3 KO). A significant reduction in the fertilization rate was observed for *Dot1l*-KO indicating that *Dot1l*-KO sperm are less functional than CTL sperm (Fig 2E). In addition to *in vivo* fertility monitoring, we performed *in vitro* fertilization (IVF) assays using the same number of spermatozoa per experiment. Using spermatozoa from *Dot1l*-KO males, no oocyte was fertilized *in vitro*, while spermatozoa from control males (including WT, CTL and HET males) led to the fertilization of ∼37% of oocytes (Fig 2E, and see Appendix FigS2G for individual values). When using oocytes devoid of zona pellucida, in order to bypass the zona pellucida crossing step, spermatozoa from *Dot1l*-KO males still performed very poorly compared to that of CTL males (∼3 % of fertilized oocytes using *Dot1l*-KO sperm compared to ∼88 % for CTL, Fig 2E), confirming that *Dot1l*-KO dramatically reduces sperm fertilizing ability. IVF is known to be less efficient and to lead to lower fertilization rates than in vivo. It is therefore more sensitive to sperm defects, as observed in other studies (Barraud-Lange *et al*, 2020; Gadadhar *et al*, 2021;Fujihara *et al*, 2010; Kawano *et al*, 2010).

### *Dot1l*-KO disrupts spermiogenesis and leads to the production of malformed nonfunctional spermatozoa

To determine the reason why *Dot1l*-KO spermatozoa are less functional than CTL ones, we investigated their morphology and motility. Spermatozoa are highly specialized cells characterized by a small head which encompasses their very compact nucleus, a long flagellum that confers motility and almost no cytoplasm. First, we observed that *Dot1l*-KO epididymal spermatozoa displayed malformed heads and abnormal flagella (characterized by thinning of some sections in their midpiece) (Figs 3A and EV2A). Inside the flagellum, the axoneme normally contains a ring of nine outer microtubule doublets and two central microtubules (9+2 axoneme). In *Dot1l*-KO sperm, ultrastructure analyses revealed abnormal axonemes, characterized by disorganized microtubules (<9+2 microtubules) (Fig 3B). Nuclear compaction was also found impaired, observed as less contrasted nuclei in KO in comparison with CTL sperm. *Dot1l*-KO sperm also showed an increased cytoplasmic retention (Figs 3B and EV2A). Using CASA (computer-assisted sperm analyses), we observed that *Dot1l*-KO sperm motility was impaired, with a reduced proportion of motile and progressively motile (i.e. swimming in an overall straight line or in very large circles) spermatozoa (Figs 3C and EV2B). This was not due to cell death as sperm vitality was comparable between CTL and *Dot1l*-KO mice (Appendix FigS3A). Axoneme/flagellum organization, nucleus compaction and cytoplasm elimination occur during the differentiation of spermatids in spermatozoa (i.e. spermiogenesis). We measured apoptosis during this transition by TUNEL assay on testicular sections, and found a higher incidence of apoptotic elongating/condensing spermatids in *Dot1l*-KO than in CTL, another indication of a defective spermiogenesis (i.e. 6% of tubules containing apoptotic elongating/condensing spermatids in CTL vs 40% of tubules in DOT1L-KO, p =0.0007, Fig 3D). Collectively, our data indicate that *Dot1l*-KO spermiogenesis is impaired at multiple levels and produces less functional spermatozoa.

**Figure 3.**
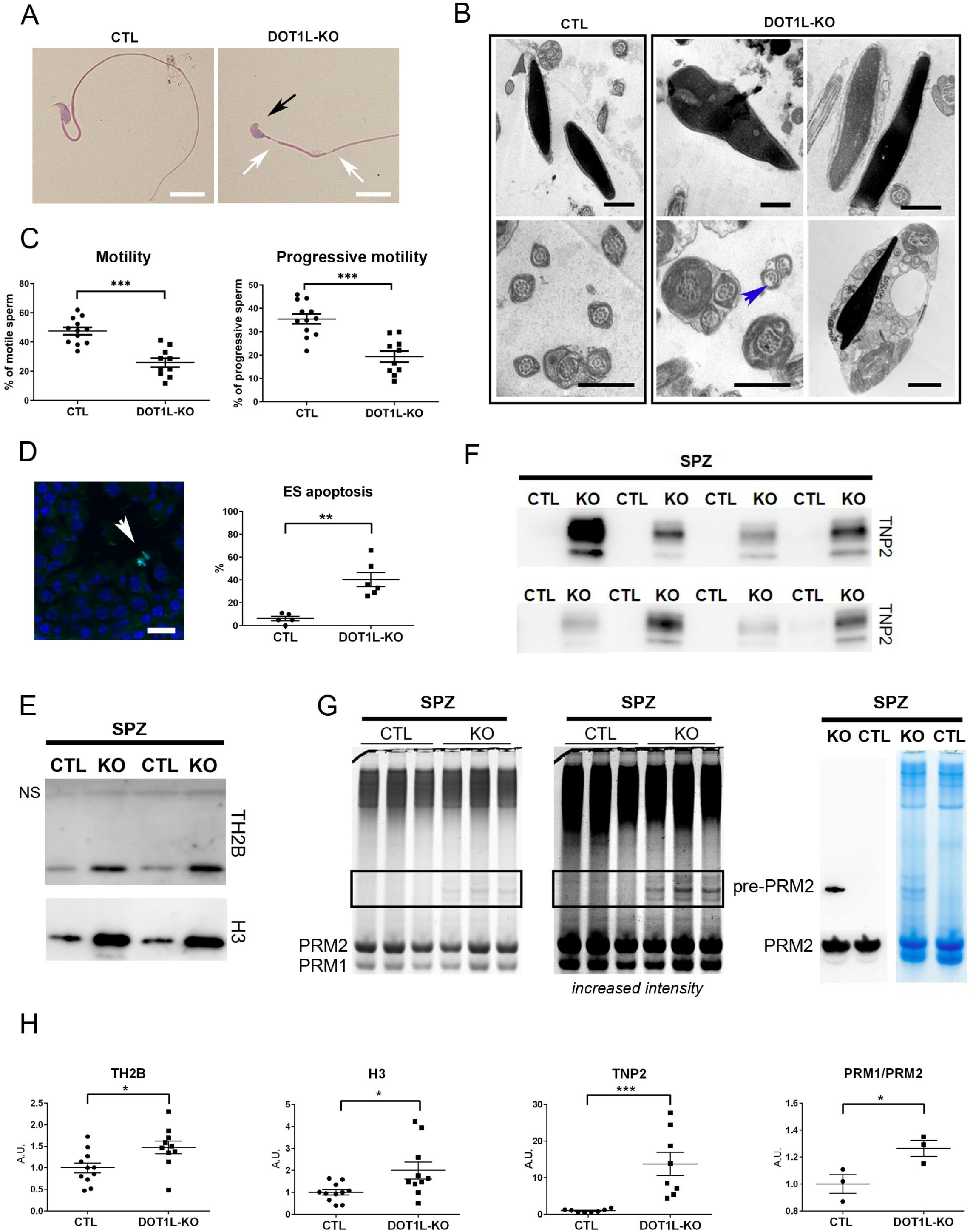
*Dot1l*-KO in male germ cells leads to multiple spermiogenesis defects including flagellar abnormalities and incomplete sperm chromatin reorganization and compaction. A) Representative pictures of *Dot1l*-KO and CTL epididymal spermatozoa. The black arrow indicates an abnormal sperm head (without apical hook) and white arrows indicate thinning of the flagellum. Scale bars represent 10µm. B) Ultrastructure pictures (electron microscopy) from epididymal spermatozoa showing multiple abnormalities in *Dot1l*-KO compared to CTL: disorganized microtubules (blue arrow), impaired nuclear compaction (top panels) and increased cytoplasmic retention (bottom right panel). Scale bars represent 1µm. C) Scatter plots (mean values following angular transformation +/- standard deviation) of the percentage of motile and progressively motile spermatozoa in *Dot1l*-KO and CTL adult mice (N= 12 for CTL and 10 for *Dot1l*-KO) obtained following CASA (computer-assisted sperm analysis). Three stars indicate a p-value <0.0005 (Mann Whitney test performed after angular transformation of the percentages). D) Scatter plots showing the percentage of testicular tubules with TUNEL-positive elongating/condensing spermatids (ES) (mean per animal + standard error of the mean, N=5 for CTL and 6 for *Dot1l*-KO). Two stars indicate a p-value <0.005 (unpaired t-test performed after angular transformation of percentages). A representative picture of *Dot1l*-KO testicular sections following TUNEL assay is shown on the left. TUNEL positive elongating/condensing spermatids are visible inside the tubule (in green, white arrowhead). DAPI (blue) was used to stain nuclei. Scale bar indicates 20µm. E) Western blot detection of histones H3 and TH2B in spermatozoa from *Dot1l*-KO and CTL males. The same quantity of material was loaded in each well (i.e. extracts from 2 million spermatozoa). NS indicates a non-specific band also observed on the membrane stained with Ponceau (see Appendix FigS3C). F) Western blot detection of transition protein 2 (TNP2) in spermatozoa from *Dot1l*-KO and CTL males. The same quantity of material was loaded in each well (i.e. extracts from 2 million spermatozoa). G) Coomassie-stained protamine extracts from CTL and *Dot1l*-KO spermatozoa following acid urea gel electrophoresis (same gel, two different intensities). The same quantity of material has been loaded in each well (i.e. extracts from 1.4 million spermatozoa). Protamine 1 and 2 bands (PRM1 and PRM2, respectively) are detected at the bottom of the gel. The rectangle highlights the bands that are likely immature forms of Protamine 2 (i.e. non-cleaved precursor forms). The right panel shows a western blot detection using anti-PRM2 antibody which confirms that one of the high molecular weight band only observed in *Dot1l*-KO spermatozoa is an immature form of Protamine 2 (Pre-PRM2). The corresponding coomassie-stained gel is also shown. H) Quantification of histones TH2B and H3 (N= 11 CTL and 10 KO), of TNP2 (N= 8 CTL and 8 KO) and of PRM1/PRM2 ratio (N= 3 CTL and 3 KO) in *Dot1l*-KO and CTL spermatozoa, related to Fig 3E-G. The data shown are normalized to control mean values (+ standard error of the mean). One star indicates a p-value <0.05, and three stars, a p-value <0.0005 (unpaired t-tests). A.U. = arbitrary units. See also Fig EV2 and Appendix FigS3.

### *Dot1l*-KO modifies the chromatin of postmeiotic male germ cells, which results in abnormal chromatin reorganization and compaction in spermatozoa

During spermiogenesis, spermatid chromatin undergoes specific reorganization and compaction as most histones are removed and replaced with protamines (Bao & Bedford, 2016; Rathke *et al*., 2014). Histone variants (such as TH2B or H2AL2) and transition proteins (TNP) are highly expressed in spermatids during histone-to-protamine transition and are essential for the correct reorganization and compaction of the sperm genome with protamines (Barral *et al*., 2017; Montellier *et al*, 2011). In addition to the nuclear compaction defects observed by electron microscopy (Fig 3B), we noticed that *Dot1l*-KO sperm chromatin is more sensitive to nucleoplasmin-induced decompaction than CTL sperm chromatin. Nucleoplasmin, or NPM, is a histone chaperone known to facilitate paternal chromatin decompaction and reorganization after fertilization owing to its ability to bind histones and sperm nuclear basic proteins (Frehlick *et al*, 2007). Following NPM treatment, a higher proportion of soluble histones was found in *Dot1l*-KO compared to CTL sperm (Appendix FigS3B) in agreement with a chromatin compaction defect. We next quantified TH2B and H3 histones in epididymal spermatozoa by western blot and, despite inter individual variability, we found significantly more residual histones in *Dot1l*-KO than in CTL sperm chromatin (Figs 3E, 3H and Appendix FigS3C-D). We also quantified transition protein 2 (TNP2) and found a clear retention of this protein in *Dot1l*-KO spermatozoa while nothing could be detected in CTL sperm (Figs 3F, 3H and Appendix FigS3E). Finally, we quantified protamines on acid-urea gels and observed an accumulation of immature (non-cleaved) forms of Protamine 2 in *Dot1l*-KO spermatozoa vs. CTL (Fig 3G). A slight but significant distorted Protamine1/Protamine2 (PRM1/PRM2) ratio was also observed in *Dot1l*-KO spermatozoa (Figs 3G-3H and Appendix FigS3F).

The transition from nucleosomes to protamine-based chromatin starts in elongating spermatids with the incorporation of histone post-translational modifications (PTMs) and histone variants [for reviews see (Bao & Bedford, 2016; Oliva, 2006; Rathke *et al*., 2014)]. To better understand how DOT1L impacts on chromatin organization prior to histone removal, we quantified histone PTMs and variants by liquid chromatography-tandem mass spectrometry (LC-MS/MS) in whole testes and in elongating/condensing spermatids. In whole testis histone extracts we observed, as expected, a clear decrease of H3K79 mono- and di-methylation in *Dot1l*-KO samples compared to CTL (Appendix FigS4A). The tri-methylation of H3K79 was not detectable with this analysis, possibly due to its small stoichiometry. No other significant differences in histone H3 PTMs were detected between *Dot1l*-KO and CTL whole testicular histone extracts. However, a significant decrease in histone H4 hyper-acetylated form (i.e. H4K5ac,K8ac,K12ac,K16ac) was observed (Appendix FigS4A). In elongating/condensing spermatids (ES), the stage during which chromatin is extensively remodelled, a significant decrease in H3K79 mono- and di-methylation levels was also observed (Fig 4A). Here also, H3K79 tri-methylation was not detectable. Several other quantitative changes in histone PTMs were observed such as modest increases in H3K23ac, H2AR88me1 and TH2BK35ac but also a dramatic decrease of H4K20 or R23 mono-, di- and tri-methylated forms (Fig 4 and Appendix FigS4B). Quantification of histone variants showed significant changes in H1 variants and in canonical H2A (Appendix FigS4C). The most remarkable change was, as in whole testes, a severe decrease in H4 acetylation in *Dot1l*-KO elongating/condensing spermatids, in particular of hyper-acetylated forms such as H4K5ac,K12ac,K16ac, H4K8ac,K12ac,K16ac and H4K5ac,K8ac,K12ac,K16ac (Fig 4B). In elongating spermatids, histone H4 hyper-acetylation is an essential step of histone-to-protamine transition, as it facilitates chromatin loosening and nucleosome disassembly prior to histone removal [for review see (Oliva, 2006)]. By western blot and immmunofluorescence, we also observed a decrease in H4 acetylation in *Dot1l*-KO elongating/condensing spermatids compared to CTL using anti-H4K16ac or anti-poly H4ac antibody (Fig EV3 and Appendix FigS4D).

**Figure 4.**
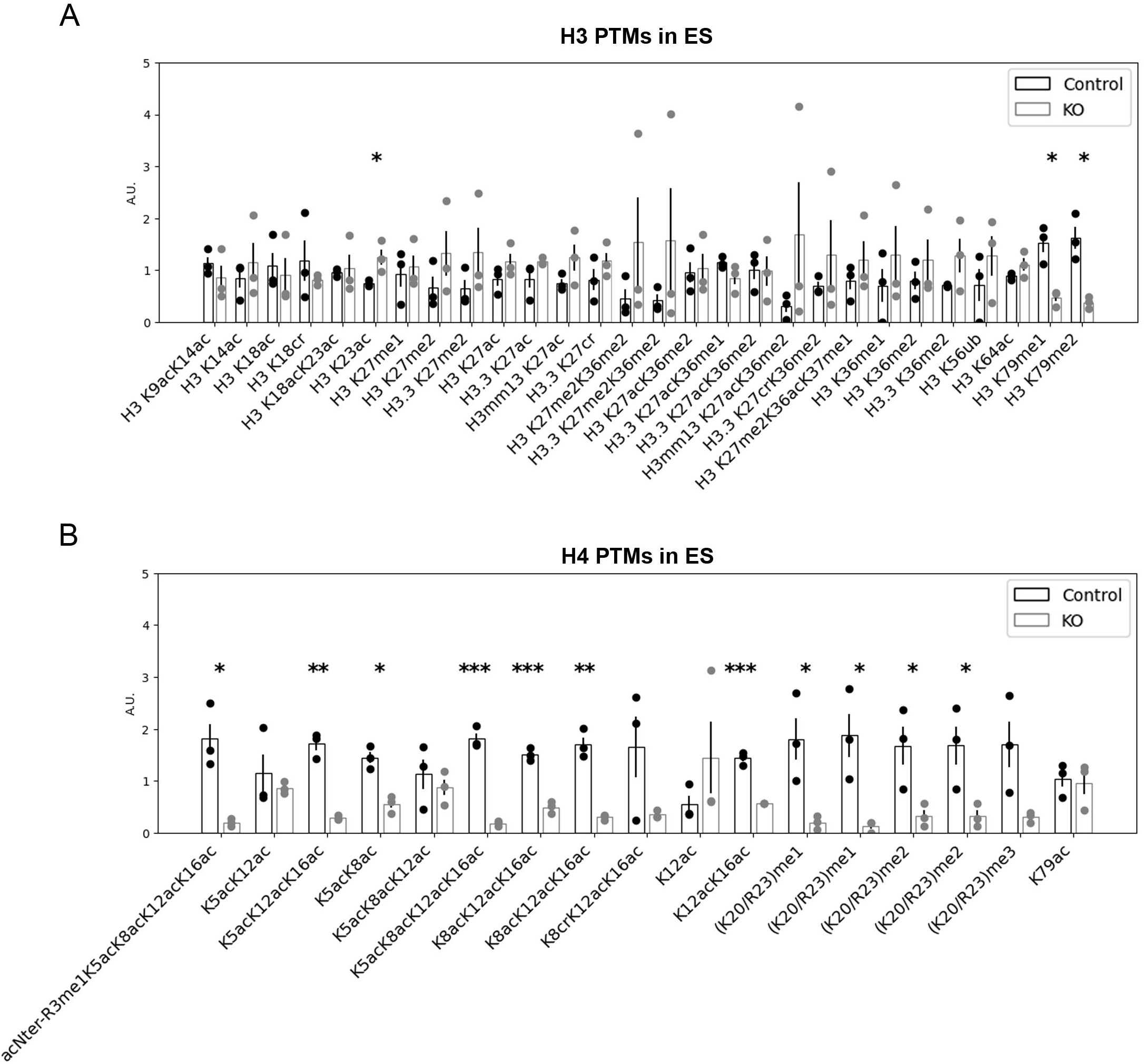
*Dot1l*-KO modifies the chromatin of elongating/condensing spermatids. Bar plots showing the quantification of post-translational modifications (PTMs) in histones H3 and H4 in ES (elongating/condensing spermatids) by LC-MS/MS. After normalization to be at constant amount of histone H3 or H4 in each analyzed sample (see Material and methods), mass spectrometry signals were divided by the average signal in both conditions (CTL and KO), so as to be able to represent all peptides in the same figure, whatever their MS intensity. Quantitative data obtained on individual biological replicates were plotted as dots, and the height of the bars indicates the average value over technical replicates. T-tests were performed to estimate the possible significant difference of modified peptide abundances between the two genotypes. One star indicates a p-value <0.05, two stars a p-value<0.005 and three stars a p-value <0.0005. When interpreting our mass spectrometry data against the database MS_histoneDB, we were able to identify various H3 variants, namely canonical H3 and variants H3.3, testis-specific H3t and H3mm13. “H3” indicates sequences shared among several variants; “H3.3”, “H3t” or “H3mm13”, each variant. A.U. = arbitrary units. See also Figs EV3, 4 and Appendix FigS4.

We next quantified histone PTMs and variants at an earlier stage of spermiogenesis, in round spermatids (RS), and did not see any decrease in H4 acetylation or H4K20 methylation in KO vs. CTL RS (Fig EV4) in contrast with what was observed in ES (Fig 4). Apart from the expected decrease in H3K79 methylation, the only quantitative changes that were detected in RS were increases in H3K18 crotonylation (H3K18cr), in H4K8cr,K12ac,K16ac and in H1 variants (Fig EV4 and Appendix FigS4C,E). In spermatozoa, some H3K79 methylation has been detected on persistent histones (Luense *et al*, 2016; Moretti *et al*., 2017) and a significant decrease in H3K79me1 and H3K79me2 levels was visible in *Dot1l-* KO spermatozoa using LC-MS/MS. Apart from a modest increase in H3K18ac, no other changes were detected at this stage (Appendix FigS4F).

All these observations indicate that *Dot1l* KO has a profound impact on postmeiotic chromatin in ES at the time of its reorganization, which disrupts histone-to-protamine transition and leads to a higher proportion of retained histones, transition proteins and unprocessed protamine 2 in spermatozoa.

### DOT1L regulates the expression of genes involved in essential processes of spermatid differentiation

In order to gain insight into the gene expression changes associated with *Dot1l* deficiency in male germ cells, we performed RNA-Seq analyses on purified germ cells from 3 different spermatogenic stages, each in 5 replicates: primary spermatocytes (SC), secondary spermatocytes (SCII) and round spermatids (RS) (Fig 5A). We did not include elongating/condensing spermatids in this analysis because their transcriptome is very similar to that of round spermatids, and, at this stage, transcription progressively shuts down as a consequence of histone-to-protamine transition and genome compaction. RNA-Seq differential analyses showed hundreds of significantly deregulated genes in *Dot1l*-KO vs CTL samples (DEseq2, with a Fold change > 1.5 and FDR < 0.05). In parallel, we performed a less stringent analysis (DEseq2, with a Fold change > 1.5 and p < 0.05, then *a posteriori* validation of FDR) and found between ∼600 and 1000 deregulated genes at each stage (Fig 5B and Appendix FigS5A). With both analyses, the number of deregulated genes increased with progression of male germ cell differentiation and was therefore highest in RS (Fig 5B and Appendix FigS5A), and approximately half of deregulated genes were found in common in at least 2 stages (Appendix FigS5A). In the three cell types, more genes were found upregulated than downregulated in KO samples (Fig 5B) in agreement with results obtained recently in *Dot1l*-KO somatic cells (Aslam *et al*, 2021; Cattaneo *et al*, 2022). To link gene deregulation in *Dot1l-*KO mice with DOT1L methyltransferase activity, we compared deregulated genes with the genomic location of H3K79me2 that we obtained by ChIP-seq in RS. H3K79me2 is the best characterized H3K79 methylation mark and is known to mark the body of transcriptionally active genes. H3K79me1 and me3 have similar trends, though broader for the former, and more restricted for the latter (Vlaming & van Leeuwen, 2016). Here, in wild type RS, H3K79me2 was found enriched at the body (with the strongest signal found near the Transcriptional Start Site) of genes which were downregulated in the KO but not those found upregulated (Fig 5C and Appendix FigS5B). Observation of ChIP peaks confirmed the presence of H3K79me2 mark at down but not upregulated genes (Fig 5D).

**Figure 5.**
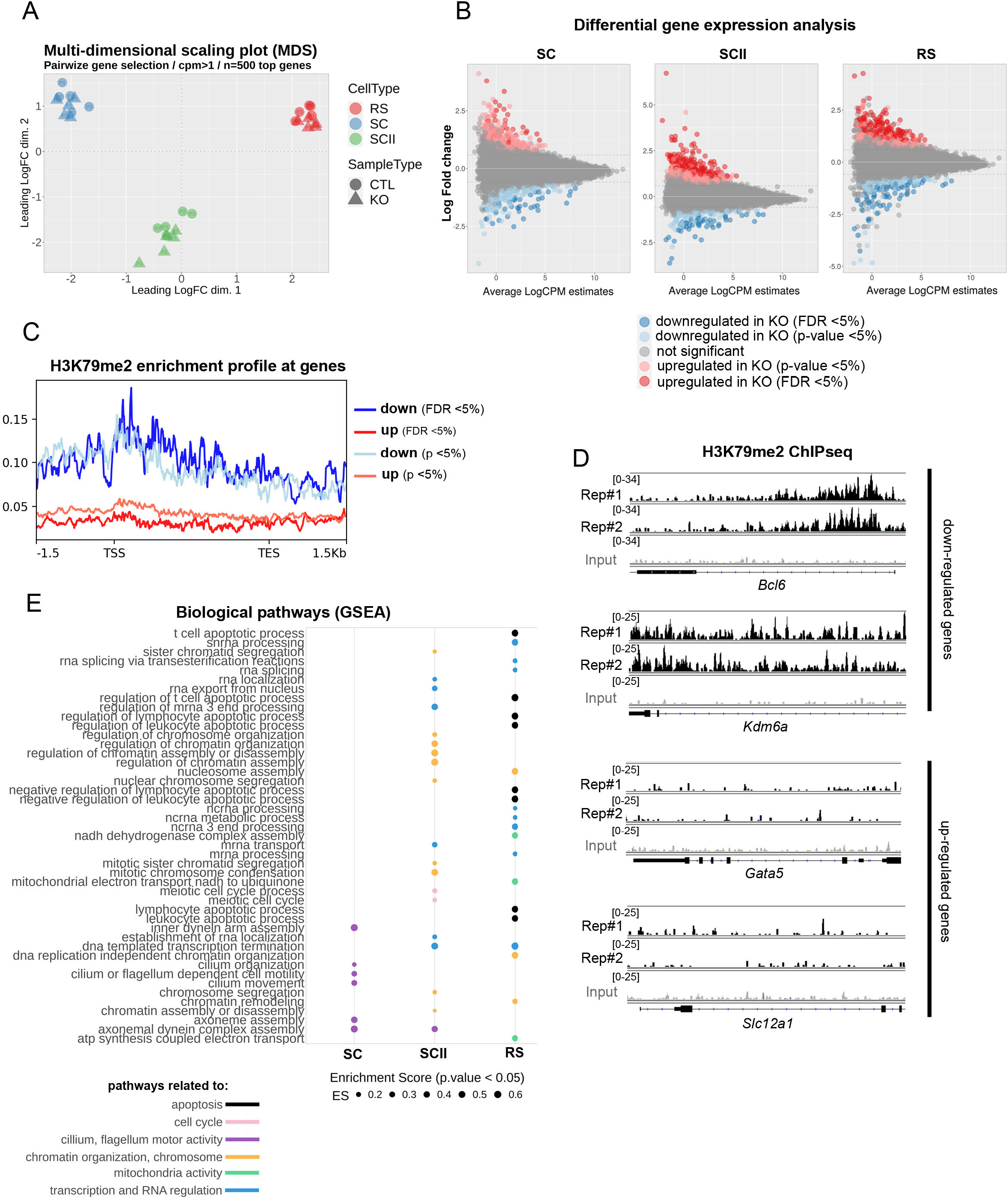
*Dot1l*-KO leads to the deregulation of genes involved in flagellum function, transcription regulation, chromatin organization, mitochondria function and apoptosis. A) Multi-dimensional scaling plot on the 500 top genes in *Dot1l*-KO vs. CTLs primary spermatocytes (SC), secondary spermatocytes (SCII) and round spermatids (RS) using pairwise gene selection. B) MD plot of the genes found deregulated in *Dot1l*-KO vs. CTLs SC, SCII and RS, using differential gene expression analyses with two different parameters (FDR<0.05 and p-value<0.05). C) H3K79me2 enrichment profile at deregulated genes (same parameters as in B). D) H3K79me2 enrichment profile at selected deregulated genes, in two biological replicates of ChIP-Seq performed in wild type RS (Rep#1 and 2. The input value is shown in grey. E) GSEA analysis of RNA-Seq data from *Dot1l*-KO vs CTL SC, SCII and RS. The figure shows all the biological pathways found significantly downregulated in *Dot1l*-KO primary spermatocytes (SC), secondary spermatocytes (SCII) and round spermatids (p<0.05), ranked by their enrichment score (ES). See also Fig EV5 and Appendix FigS5.

We next investigated deregulated pathways using GSEA (Gene Set Enrichment Analysis), a threshold-free method which considers all genes based on their differential expression rank, and found a downregulation of biological pathways related to apoptosis, transcription/RNA regulation, chromatin/chromosome organization and mitochondria activity in *Dot1l*-KO vs. CTL round spermatids (Fig 5E). Prior to spermiogenesis, pathways related to cilium or flagellum motor activity were found downregulated in *Dot1l*-KO primary spermatocytes while, in *Dot1l*-KO secondary spermatocytes, pathways related to chromatin/chromosome organization were the most significantly deregulated (Fig 5E). When looking at gene ontology for “cellular components”, pathways related to ribosomal structure or to mitochondrial proteins were found deregulated in RS (Fig EV5). Strikingly, several *Slc* genes were found deregulated in *Dot1l*-KO RS (Appendix FigS5C). *Slc* genes encode solute carriers, and several of them have been implicated in sperm motility *via* their effects on flagellum differentiation and/or sperm energy production (Kuang *et al*, 2021; Maruyama *et al*, 2016; Toure, 2019).

In agreement with the downregulation of pathways related to apoptosis, *Bcl6, Jak3* and *Tsc22d3* which encode proteins with anti-apoptotic effects (Aguilar *et al*, 2014; Kurosu *et al*, 2003; Thomis *et al*, 1997) were found significantly downregulated in RS. *Blc6* codes for a transcription repressor (Chang *et al*, 1996) and appears to be a direct target of DOT1L since it is downregulated upon *Dot1l* loss, and enriched in H3K79me2 in wild-type RS (Fig 5E). Interestingly, an unsupervised search for DNA motifs identified BCL6/BCL6B DNA binding site as the most significantly enriched motif in the promoter of upregulated genes (15 upregulated genes, Appendix FigS5D). This finding is in agreement with a model in which downregulated genes are direct H3K79me2/DOT1L targets while upregulated genes are indirect targets.

## Discussion

In the present study, using a conditional knockout mouse model, we demonstrate that the H3K79 methyltransferase DOT1L is essential for gene regulation and chromatin remodelling during spermatid differentiation.

DOT1L is highly conserved among eukaryotes. In mammalian it has been extensively studied and found to be involved in many different biological processes such as mixed lineage leukaemia (Nguyen *et al*, 2011; Okada *et al*, 2005), cell cycle (Kim *et al*., 2014), development (Jones *et al*, 2008), reprogramming (Onder *et al*, 2012), DNA damage repair (Lin *et al*, 2009; Zhu *et al*, 2018) or transcription activation (Steger *et al*, 2008; Wang *et al*, 2008). In the male germline, DOT1L is highly expressed at the RNA and protein levels, in particular in postmeiotic cells (Dottermusch-Heidel *et al*., 2014a; Dottermusch-Heidel *et al*., 2014b; Moretti *et al*., 2017). Before meiosis, DOTL1 has been found to be required for the self-renewal of adult stem cells. Indeed, with a *Dot1l* KO induced by a Cre recombinase expressed earlier than the one used in the present study, Lin et al. observed a progressive (age-dependent) loss of all male germ cell types, ultimately resulting in a Sertoli cell only syndrome (i.e. absence of germ cells) (Lin *et al*., 2022). In the *Dot1l*-KO males we characterized here, we also found empty tubules as well as a decrease of all germ cell types, but milder, which enabled us to address the role of DOT1L in adult postmeiotic germ cells (i.e. spermatids). Strikingly, we found that *Dot1l*-KO spermatozoa present multiple anomalies such as thinner and distorted flagella, cytoplasmic retention and impaired nuclear compaction. As a result, KO spermatozoa are less motile and their fertilizing ability is compromised. These defects are associated with the deregulation of hundreds of genes in *Dot1l*-KO meiotic and postmeiotic germ cells. Previous studies have shown that spermatid differentiation is associated with the specific expression (or highly enriched expression) of thousands of genes, and that this genetic program starts as early as in primary spermatocytes (Chen *et al*., 2018; da Cruz *et al*., 2016; Ernst *et al*., 2019; Green *et al*., 2018; Soumillon *et al*., 2013). Here, our analyses indicate that pathways related to “cilium/flagellum assembly or motility” are deregulated from the spermatocyte stage, and those related to “nucleosome assembly, chromatin remodelling” are deregulated in secondary spermatocytes and round spermatids. A significant proportion of *Dot1l*-KO spermatids undergo apoptosis and this was also visible by RNA-Seq with the deregulation of “apoptosis” related pathways in RS, including the downregulation of genes with anti-apoptotic effects such as *Bcl6* or *Jak3* (Kurosu *et al*., 2003; Thomis *et al*., 1997). Interestingly, several potential BCL6 target genes were identified among the genes which are upregulated in *Dot1l*-KO spermatids. Like other upregulated genes, most of them are devoid of H3K79me2 in wild type spermatids and may not be directly regulated by DOT1L. In contrast, *Bcl6* and many other downregulated genes in *Dot1l*-KO spermatids are marked by H3K79me2 in wild-type cells and are likely directly regulated by DOT1L. Our findings are in agreement with recent studies in which *Dot1l* knockout was investigated in B cells, T cells or cardiomyocytes. Each time, H3K79me2 was found to be enriched at the body of downregulated genes but not of upregulated genes (Aslam *et al*., 2021; Cattaneo *et al*., 2022; Kwesi-Maliepaard *et al*, 2020).

Strikingly, many genes encoding members of the solute carrier family were deregulated in *Dot1l*-KO male germ cells. Several members of this family such as *Slc22a14* and *Slc26a8* have previously been implicated in sperm motility *via* their effects on flagellar differentiation and/or sperm energy production (Kuang *et al*., 2021; Maruyama *et al*., 2016; Toure, 2019); the deregulation of *Slc* genes with similar function could therefore contribute to the sperm flagellar and motility defects of *Dot1l-* KO males. Overall, these data indicate that *Dot1l*-KO-induced gene deregulation is responsible for the multiple defects of postmeiotic male germ cell differentiation.

In addition to being involved in the regulation of spermatid gene expression, we show that DOT1L is essential for spermatid chromatin remodelling and subsequent reorganization and compaction of spermatozoon genome. Indeed, the chromatin of *Dot1l*-KO spermatozoa is less compact, and characterized by a higher level of retained histones, of transition protein 2 and of immature forms of protamine 2, as well as altered protamine 1/2 ratio. Several other studies have observed similar anomalies in particular retained histones and immature forms of protamine 2, as a result of incomplete histone-to-protamine transition, such as *Tnp1-*KO (*transition protein 1*) (Yu *et al*, 2000), *H2al2*-KO (*H2A histone family member L2A*) (Barral *et al*., 2017), deletion of the mouse Y chromosome long arm (Yamauchi *et al*, 2010), or *Kat2a* –KO (a.k.a. *Gcn5)* which encodes the lysine acetyltransferase 2A (Luense *et al*., 2019). Similarly, a recent study has shown that aberrant pre-PRM2 cleavage leads to altered levels of histone variants and transition proteins in sperm (Arevalo *et al*, 2022). In all those mouse models as well as in the *Dot1l*-KO presented here, abnormal sperm chromatin content is associated with fertility problems. This is also the case in humans, since impaired protamine 1/2 ratio and accumulation of pre-PRM2 are associated with male infertility (de Mateo *et al*, 2009; Torregrosa *et al*, 2006).

In elongating spermatids, histones – in particular histone H4 – are hyper-acetylated prior to their replacement. This phenomenon is a prerequisite for histone-to-protamine transition as it creates a permissive state for histone removal [for reviews see (Bao & Bedford, 2016; Oliva, 2006; Rathke *et al*., 2014)]. H3K79me2/3 levels peak at the time of H4 hyperacetylation (Dottermusch-Heidel *et al*., 2014a; Dottermusch-Heidel *et al*., 2014b; Moretti *et al*., 2017) and we show here that both H3K79 methylation and H4 hyperacetylation are drastically and specifically reduced in *Dot1l*-KO elongating spermatids. Several acetyltransferases are involved in H4 hyperacetylation such as KAT2A, CBP-P300 or NuA4 complex (Boussouar *et al*, 2014; Luense *et al*., 2019; Shiota *et al*., 2018) but none of them was found deregulated in *Dot1l*-KO spermatids. We therefore propose that the decrease in histone acetylation is a consequence of impaired H3K79 methylation. In other contexts than the male germline, H4 acetylation and H3K79 methylation have previously been shown to influence each other in different ways. In 2016, Gilan *et al*. proposed that DOT1L-mediated H3K79me2 leads to a more open chromatin which facilitates the recruitment of the histone acetyltransferase P300 which, in turn, increases histone H4 acetylation (in particular of H4K5ac), in a cell model of acute myeloid leukemia (Gilan *et al*, 2016). In contrast, H4K16ac has been shown to stabilize the binding of yeast Dot1 and human DOT1L on the nucleosome and consequently stimulates its methyltransferase activity *in vitro* (Valencia-Sanchez *et al*, 2021). In elongating spermatids, H3K79 methylation and H4 acetylation could therefore contribute to increase each other signal. It would be interesting to test if H3K79 methylation level is affected in spermatids in which histone acetylation is impaired, for instance in *Kat2a* knockout spermatids.

In our study, we also observed a decrease in the three methylation states of H4K20 in elongating spermatids. This is in line with the observation made by Jones et al. (Jones, Su et al., 2008) that H4K20 tri-methylation level is reduced at centromeres and telomeres of *Dot1l*-KO embryonic stem cells. H3K9 di-methylation (but not tri-methylation) was also found reduced in KO embryonic stem cells, but this histone PTM could not be detected in our analyses. Methylation of H4K20 has been shown to be associated with chromatin accessibility. H4K20me1, by changing the conformation of nucleosomes, increases accessibility for other factors involved in DNA processing and expression (Shoaib *et al*, 2021). In contrast, H4K20me2 and H4K20me3 are associated with chromatin compaction (Evertts *et al*, 2013). For the moment, the impact of these marks during spermatid differentiation is unknown, but H4K20me3 is one of the most dynamic histone modifications during spermatogenesis, with a peak in elongating spermatids ; H4K20me2 follows the same pattern but is less abundant than H4K20me3 (Luense *et al*., 2016; Wang *et al*, 2021). One can therefore speculate that H4K20 methylation is also important for histone to protamine transition. Interestingly, Ho *et al*. have found that the H4K20me1 methyltransferase, SET8, and DOT1L (which is a non-SET domain methyl-transferase) bind to the nucleosomal acidic patch by a similar mechanism (Ho *et al*, 2021). Further work will be needed to understand the mechanism by which DOTL1 and/or H3K79 methylation may influence H4K20 methylation.

Other, more modest quantitative changes in histone PTMs and variants were observed in elongating spermatids. They could be compensatory mechanisms for H3K79 methylation or H4 acetylation decrease, as observed in *Th2b* knock-out spermatids where H2B was upregulated and several histone PTMs changed to compensate for the loss of H2B variant TH2B (Montellier *et al*., 2013). The deregulation of genes associated to “chromatin function” could also contribute to the changes in chromatin content observed in *Dot1l*-KO spermatids.

In conclusion, our data show that DOT1L is indispensable for the differentiation of postmeiotic germ cells into functional spermatozoa. Indeed, DOT1L regulates the expression of genes essential for spermatid differentiation and, DOT1L-mediated H3K79 methylation is necessary for proper histone H4 hyperacetylation in elongating spermatids and for an efficient histone-to-protamine transition and subsequent compaction of the sperm chromatin.

## Materials and methods

### Mouse strains and genotyping

Conditional *Dot1l* knockout mice were obtained from the Knockout Mouse Project (CSD29070) (to F.V.D). This line has previously been described in (Jo *et al*., 2011). In these mice, *Dot1l* exon 2 is flanked by loxP sites (see Appendix FigS1D). Recombination of loxP sites by Cre recombinase leads to the deletion of exon 2 and consequently a frameshift and premature stop codon in *Dot1l* Coding DNA sequence. Tg(*Stra8-iCre*)1Reb/J were obtained from the Jackson lab (JAX stock #017490) (Sadate-Ngatchou *et al*., 2008). All animals analyzed in this study were from a C57BL6/J background. Genotypes of animals were determined by polymerase chain reaction (PCR) using *Dot1l* primers F+R1+R2 (Appendix FigS1D) or *iCre* primers as described on the Jackson lab website. The latter PCR was performed with *Ymtx* internal control (see Appendix Table S1 for primer list and sequences).

Unless specified, all experiments were performed on (adult) 3 to 5 month-old males. Reproductive parameters were analyzed on two types of *Dot1l* knocked-out animal models. First, on animals hosted in a conventional animal house (cah) and of the following genotypes: *Dot1l*^*Fl/Fl*^ ; *Stra8-Cre* mice (DOT1Lcah) and *Dot1l*^*Fl/Fl*^ siblings (without *Stra8*-Cre transgene) as controls (CTLcah). Second, on animals hosted in a Specific Pathogen-Free (SPF) animal house and of the following genotypes: *Dot1l*^*Fl/Δ*^ ; *Stra8-Cre* (hereafter called DOT1L-KO) and *Dot1l*^*Fl/Fl*^ siblings as controls (hereafter called CTL). Analyses of the reproductive parameters of both types of KO males (*Dot1l*^*Fl/Fl*^;*Stra8-Cre* or *Dot1l*^*Fl/Δ*^; *Stra8-Cre*) gave similar results (Appendix FigS2A and B). Heterozygous *Dot1l*^*Fl/Δ*^ males (HET) were also analyzed. They did not differ from controls (Appendix FigS2A-E). *Dot1l*^*Fl/Δ*^; *Stra8-Cre* animals were generated because it had been shown that *Stra8-iCre* is not 100 % efficient (Bao *et al*., 2013). In these animals, one allele of *Dot1l* is floxed and one allele is already deleted (Δ) to increase the efficiency of floxed exon excision upon *Cre* recombinase expression. Nevertheless, some remaining DOTL1 protein signal was observed by Western blot and Immunofluorescence (see Figs 1B, D and Appendix FigS1E-I). All other phenotypic analyses and all molecular analyses were performed on *Dot1l*^*Fl/Δ*^ ; *Stra8-Cre*, hereafter called *Dot1l*-KO, and on *Dot1l*^*Fl/Fl*^ siblings as CTL. Mice from mTmG line (Muzumdar *et al*, 2007) were crossed to *Stra8-Cre* line to verify *Stra8-Cre* recombinase location and activity.

The mice were fed ad libitum with a standard diet and maintained in a temperature- and light-controlled room. Animal procedures were approved by Universite de Paris ethical committee (Comite d’Ethique pour l’Experimentation Animale; registration number CEEA34.JC.114.12, APAFIS 14214-2017072510448522v26).

### Fertility tests

*Dot1l*-KO and CTL males, aged from 8 to 10 weeks, were housed with two wild-type C57BL6/J females per cage (aged from 6 weeks, Janvier Labs, France) for up to 4 months. Vaginal plugs were checked and females separated when found. For each group, litter size and number were assessed.

Another series of test were performed too evaluate the fertilization rate following mating (in vivo fertilization). For this WT females were first superovulated by injection of 5 IU of PMSG (pregnant mare’s serum gonadotrophin; Intervet, France) followed by 5 IU of hCG (human chorionic gonadotrophin; Intervet) 48 h later. Females were then mated with WT or *Dot1l-*KO males at night and the next day females with a vaginal plug were isolated. Oocytes were collected from ampullae of oviducts and freed from the cumulus cells by brief incubation at 37°C with hyaluronidase (Sigma, St. Louis, MO, USA) in M2 medium (Sigma). Oocytes were rinsed and the number of fertilized oocytes was evaluated using DAPI immune-fluorescent staining to visualize the DNA.

### IVF

Oocyte preparation: WT C57BL6/J female mice aged 6 to 8 weeks (Janvier Labs, France) were superovulated with 5 IU of pregnant mare serum gonadotropin (PMSG) and 5 IU human chorionic gonadotropin (hCG) (Intervet, France) 48 hours apart. About 14 hours after hCG injection, animals were sacrificed by cervical dislocation. Cumulus oocyte complexes were collected by tearing the ampulla wall of the oviduct, placed in Ferticult medium (FertiPro N.V, Belgium) supplemented with 3 % BSA (Sigma–Aldrich), and maintained at 37°C under 5 % CO2 in air under mineral oil (FertiPro N.V, Belgium). When experiments were performed with Zona-free oocytes, cumulus cells were first removed by a brief exposure to hyaluronidase IV-S (1 mg/ml, Sigma–Aldrich). The zona pellucida was then dissolved with acidic Tyrode’s (AT) solution (pH 2.5) (Sigma–Aldrich) under visual monitoring. Zona-free eggs were rapidly washed five times and kept at 37°C under 5 % CO2 in air for 2 to 3 hours to recover their fertilization ability.

Capacitated sperm preparation: mouse spermatozoa were obtained from the cauda epididymides of DOT1L-KO, CTL, HET and WT C57BL6/J males (aged 8 to 12 weeks) and capacitated at 37°C under 5 % CO2 for 90 minutes in a 500 µl drop of Ferticult medium supplemented with 3 % BSA, under mineral oil.

*In vitro* fertilization: cumulus-intact and Zona-free eggs were inseminated with capacitated spermatozoa for 3 hours in a 100 µl drop of Ferticult medium, 3 % BSA at a final concentration of 10^6^ or 10^5^ per ml, respectively. Then, they were washed and directly mounted in Vectashield/DAPI (Vector laboratories, CA, USA) for observation under UV light (Nikon Eclipse E600 microscope). Only oocytes showing at least one fluorescent decondensed sperm head within their cytoplasm were considered fertilized.

### Testicular germ cell purification by elutriation or FACS

Germ cells were purified from adult males using fluorescence-activated cell sorting (FACS) or elutriation as previously described in (Cocquet *et al*, 2009; Comptour *et al*, 2014) and more recently in (Crespo *et al*, 2020). Elutriated fractions with a purity ∼95–99% for elongating/condensing spermatids and ∼80% for round spermatids, were used for LC-MS/MS analyses. Highly enriched fractions of primary or secondary spermatocytes (>90%) and round spermatids (∼99%) were used for RNA-Seq analyses. Flow cytometric analysis of testicular cell suspension were performed as previously described (Corbineau *et al*, 2017; Ragazzini *et al*, 2019).

### Sperm collection and purification with Percoll

Spermatozoa were extracted from cauda epididymis in pre-warmed (37°C) M2 medium (Sigma-Aldrich) by gentle pressure. Then, epididymides were perforated with a thin needle and incubated in M2 for 10 min at 37°C to allow remaining sperm to swim up. For molecular analyses, spermatozoa were next purified on a Percoll gradient. In brief, spermatozoa were centrifuged at 2000 g for 10 min and pellets incubated in somatic cell lysis buffer (1X PBS, 0.1% SDS, 0.5% Triton X-100) for 10 min on ice. Sperm cells were washed with PBS-BSA (1X PBS, 0.5% BSA, 2mM EDTA) and briefly sonicated to remove flagella (ON 5 sec – OFF 30 sec x 3 Cycles, bioruptor Pico, Diagenode). The samples were transferred in low retention tubes, loaded on 50% Percoll (Sigma-Aldrich) and centrifuged at 2500 g for 5 min to remove somatic cells and flagella. This step was repeated once and sperm cells were washed in PBS-BSA twice and counted (minimum 1 000 cells). The purity was >99.7%. Spermhead pellets were snap-frozen in liquid nitrogen and stored at -80°C prior to use.

### Nucleoplasmin decompaction

Experiments were performed as previously described in (Yamaguchi *et al*, 2018). Following Percoll purification, sperm heads were washed in 1X PBS, 0.5% Tween, permeabilized with 0.02 U/M Streptolysin O (Sigma-Aldrich) in 1X PBS for 10 min on ice and washed once with 1X PBS, 0.5% Tween. Cells were then incubated with 1X PBS containing 1 mM DTT for 10 min at 37°C, washed with 1X PBS, 0.5% Tween, resuspended in KH buffer (100 mM KCl, 20 mM Hepes-KOH, pH 7.7) in low retention tubes and stored at -80°C for later use. Ten million of sperm heads were resuspended in 1 mL NPM treatment buffer (20 mM HEPES-KOH, pH 7.7, 100 mM KCl, 2.5 mM MgCl2, 5 mM Na-But, 10 mM DTT, 1X complete EDTA-free protease inhibitors, 250 µM NPM) and incubated for 2 h at 37 °C in a Thermomixer (Mixer HC, Starlab) at 1000 rpm. After centrifugation at 20 000 g for 5 min at 4°C, cells were washed with 5 mM Na-But in 1X PBS, 0.5% Tween twice and sperm heads were fixed with 1% PFA for 10 min at RT, quenched with 250 mM glycine and washed in 5 mM Na-But in PBS/Tween 0.5% twice.

After fixation, sperm pellets were resuspended in NP40 Buffer (10 mM Tris-HCl, pH 8.0, 10 mM NaCl, 0.5% NP40, 1X complete EDTA-free protease inhibitors, Sigma Aldrich) and incubated on ice for 30 min. Samples were washed with 1X PBS, 0.5% Tween, 5 mM Na-But, and resuspended in 200 µL of sonication buffer (10 mM Tris-HCl, pH 8.0, 20% glycerol, 0.25% SDS, 5 mM Na-But, 1X complete EDTA-free protease inhibitors). Sperm cells were sonicated (ON 30sec – OFF 30sec x13 cycles, Bioruptor Pico, Diagenode) and the sheared chromatin was centrifuged at 20 000 x g for 10 min at 4°C. Supernatants were transferred in new 1.5 mL tubes and pellets resuspended in 200 µL of KH buffer. Prior to western-Blot analyses, both supernatant and pellets were resuspended in 4X NuPage/10% β-mercapto-ethanol (ThermoFisher), heated 10 min at 95°C and briefly sonicated.

### Histone extraction from spermatozoa

Experiments were performed as described in (Crespo *et al*., 2020) with minor modifications. In brief, five million of Percoll-purified and sonicated spermheads were incubated in 50mM DTT for 30 min (at 4°C) then mixed with sulfuric acid (0.4M final volume), sonicated and acid extracted with 20% trichloroacetic acid (TCA). Histone pellets were washed with cold acetone containing 0.05% HCl, dried at room temperature, and resuspended in 50µl of SDS-PAGE loading buffer containing 10% β-mercapto-ethanol.

### Western blot

Electrophoresis was performed in polyacrylamide gels at 120 V in denaturating buffer containing 25 mM Tris Base, 190 mM glycine, 0.1 % SDS and proteins were transferred on nitrocellulose membranes (GE Healthcare). Membranes were then rinsed and incubated 3 min in Ponceau stain to visualize transfer efficiency. Then membranes were incubated in 1X PBS, 0.01 % Tween, 5 % milk. All primary antibodies were incubated over night at 4°C (see Appendix Table S2 for references and dilutions) and 2 h at room temperature for secondary antibodies. The revelation was performed with SuperSignal West Pico Plus® ECL from ThermoFisher (34580) and Immobilon ECL Ultra Western HRP substrate from Millipore (WBULS0100) on ImageQuant™ LAS 4000 imager.

### Extraction and analysis of protamines

PRM1/PRM2 protamine ratio was analyzed following procedures of protamine-rich fraction of sperm nuclear proteins extraction and analysis previously described, with minor modifications (Soler-Ventura *et al*, 2018). Briefly, histones and other basic proteins loosely attached to DNA were extracted incubating sperm in 0.5 M HCl for 10 min at 37 °C after vortexing, and centrifuged at 2000 *g* 20 min at 4 °C. This step was repeated three times. Resulting pellet was resuspended in 0.5 % Triton X-100, 20 mM Tris-HCl (pH 8), and 2 mM MgCl_2_. After centrifugation at 8940 *g* 5 min at 4 °C, the sediment was resuspsended in milliQ H_2_O with 1 mM PMSF, centrifuging again at the same conditions. Chromatin was then denaturated by resuspending the pellets in 20mM EDTA, 1mM PMSF, 100mM Tris-HCl-pH8) and adding 1 volume of 575 mM DTT in 6 M GuHCl prior vortexing. The solution was incubated at 37 °C with 0.8 % 4-vynilpyridine to inhibit cysteine disulfide bonds for 30 min, vortexing every 5 min, in fumehood and in dark conditions. Chromatin was then precipitated with a minimum of 10 min incubation with cold ethanol at -20 °C, followed by centrifugation 12880 *g* for 15 min at 4 °C. Basic nuclear proteins (protamine-enriched fraction) were extracted from DNA incubating with 0.5 M HCl at 37 °C and recovered in the supernatant after centrifugation at 17530 *g* for 10 min at 4 °C. Precipitation was carried out with 20 % TCA on ice and centrifugation using the same conditions. Protamines were washed twice with 1 % β-mercaptoethanol in acetone and dried out at room temperature. For in-gel quantification, dried purified extracts were resuspended 5.5 M urea, 20% β-mercaptoethanol, 5% acetic acid, and separated using acetic-acid urea gel electrophoresis (AU-PAGE). Gels were stained with EZBlue™ Gel Staining Reagent (#G1041, Sigma Aldrich) and optic density of the bands corresponding to mouse PRM1 and PRM2 was quantified using Quantity One 1-D analysis software (BioRad, Hercules, CA, USA) to calculate the PRM1/PRM2 ratio. PRM1/PRM2 ratios were normalized against the mean value of the control group.

For western blot detection, basic nuclear protein extracts from 2.1M spermatozoa were loaded into AU-PAGE. Proteins were then transferred towards the negative pole onto a 0.45μm pore size nitrocellulose membrane (88018, ThermoFisher) for 4h at 4°C using an acidic transfer buffer consisting of 0.9mM acetic acid. The rest of the procedure (blocking, antibody incubation and ECL detection) was similar as for other western blot experiment (see dedicated paragraph above).

### Histological analyses

Mouse testes were fixed in 4 % buffered paraformaldehyde (PFA) for minimum 4 h, before they were cut in halves and incubated overnight in Bouin (Sigma). Testes were then washed in 70 % ethanol at room temperature for 30 min, dehydrated, and embedded in paraffin. Four µm-sections were stained with periodic acid–Schiff (PAS).

### Immunohistochemistry and immunofluorescence

Mouse testes were fixed in 4 % buffered paraformaldehyde (PFA) for minimum 4 h, before they were cut in halves and incubated overnight in 4 % PFA. Testes were then washed in 70 % ethanol at room temperature for 30 min, dehydrated, and embedded in paraffin. Immunohistochemistry or immunofluorescence experiments were performed on 4 µm-sections using Novolink polymer detection system (7140-K, Leica Micro-systems) according to the manufacturer’s instructions, or following the procedure described in (Comptour *et al*., 2014) with the following modifications. Permeabilization was performed for 15 min in 1X PBS, 0.5 % Triton X-100. Blocking was performed for 30 min to 1 h at room temperature in 1X PBS, 0.1 % Tween, 1 % BSA. Primary antibody was incubated overnight at 4°C (dilution and reference of each antibody are available in Appendix Table S2). Some slides were counterstained with Hematoxylin. For immunofluorescence experiments, lectin (L-21409 or L-32459, ThermoFisher) was diluted at 1/500 in 1X PBS and incubated for 1 h at room temperature along with secondary antibodies (see (Comptour *et al*., 2014)). Lectin was used to stain the developing acrosome and determine the stage of testis tubules as described in (Ahmed & de Rooij, 2009). DAPI (in VECTASHIELD Mounting Medium, Vectorlab) was used to stain nuclei. TUNEL assay was performed on 4 µm paraffin-embedded testicular sections using In Situ Cell Death Detection Kit, Fluorescein as described by the manufacturer (Roche, Sigma-Aldrich). GFP immunofluorescence pictures were taken with a NikonE600 microscope using the Nikon Digital Sight SD-V3 camera (DS-Fi1) with the NIS software. All other immunofluorescence pictures were taken with an Olympus BX63 microscope and analysed using ImageJ 1.48v (http://imagej.nih.gov/ij/). Immunohistochemistry pictures were taken with Perkin Elmer Lamina slide scanner and analyzed using CaseViewer software.

### Papanicolaou staining of spermatozoa

Spermatozoa were collected from cauda epididymides in M2 medium (Sigma-Aldrich) and spread onto a Superfrost Plus slide (ThermoFisher). Cells were fixed in 4 % PFA for 10 min and stained using Papanicolaou staining. Briefly, the slides were washed in 95 % ethanol, incubated in Harris hematoxylin for 3 min for nucleus counterstaining, washed and stained with OG-6 dye (RAL diagnostics 05.12013, Martillac, France) and with EA-50 (RAL diagnostics 05.12019, Martillac, France). The slides were then dehydrated and mounted with permanent mounting medium. Spermatozoa were then observed with NikonE600 optic microscope and pictures were taken with the 40X objective.

### Electron microscopy

Ten to 20 million epidydimal spermatozoa were collected in 2ml of fresh M2 medium (Sigma-Aldrich) prewarmed at 37°C. After pelleting the sperm at room temperature (300g, 10 min), the pellet was fixed in 3 % glutaraldehyde for 1 h before washing twice in 1X PBS (1000g, 10 min). Samples were then dehydrated in ethanol baths and incubated in propylene oxide before inclusion in gelatin. Slices of 60 nm were cut with a Diatome and observed with a *JEOL 1011* microscope at a magnification of 1500X or 2000X.

### Computer assisted sperm motility analysis (CASA)

Immediately after collection of epididymal spermatozoa in M2 medium (Sigma-Aldrich), sperm motility was analyzed by computer-assisted sperm analysis (CASA) systems CASA CEROS II (Hamilton Thorne, Beverly, MA, USA), using a Zeiss microscope.

### Sample preparation for RNA-Seq

Total RNA from 150,000 to 600,000 cells of sorted primary spermatocytes (SCI), secondary spermatocytes (SCII) and round spermatids (RS) was extracted using the Ambion RNAqueous micro kit (ThermoFisher) following manufacturers’ instructions. For each cell type, 5 replicates were analyzed. Quantity and quality were assessed using Bioanalyzer chips (ThermoFisher). Libraries were prepared using the NEBNext Ultra II Directional RNA Library Prep Kit (New England Biolabs) according to supplier recommendations. Paired-end sequencing of 100-bp reads was then carried out on the Illumina HiSeq4000 system.

### RNA-Seq analyses

Snakemake was used for RNA-seq analyses (Koster & Rahmann, 2018)(v. 3.9.0). Adaptors were trimmed and reads with poor quality (quality < 20) were filtered out with BBduk from BBTools (v. 38.23, https://sourceforge.net/projects/bbmap/). Alignment was performed on the mouse genome build mm10 (GRCm38.p6) using STAR (Dobin & Gingeras, 2015) (v. 2.7.2d) and Gencode vM19 gene annotation GTF with following arguments : --outMultimapperOrder Random --quantMode Genecounts --sjdbOverhang 74. For each sample, the number of aligned reads was converted in cpm (count per million). Genes with an expression level of at least > 1cpm in a minimum of 2 samples were included in the analysis, to exclude genes with a very low expression. Differential expression analysis was carried out using DESeq2 (Love *et al*, 2014) and edgeR (Robinson *et al*, 2010) packages using default parameters and a design using genotype and cell type. Differentially expressed genes (FDR<5%) were obtained using glmTreat edgeR’s function which conducts a modified likelihood ratio test (LRT) against the fold change threshold (> 1.5). This method is based on TREAT method (McCarthy & Smyth, 2009).

Two analyses were performed in parallel: one selecting the deregulated genes with a FDR < 5% (adjusted p-values), and one selecting deregulated genes with a p-value < 5% (non-adjusted p-values). On this second category of genes, the FDR was calculated *a posteriori* using Benjamini-Hochberg correction (Benjamini & Hochberg, 1995) and was confirmed to be < 5%. Enrichment analysis was performed with GSEA (v. 4.0.3, c5.all.v7.1.symbols.gmt, GO cellular components, biological pathways, or all) on gene expression cpm values between KO and control samples (Subramanian *et al*, 2005) (Fig. S5E). Beforehand, mouse Ensembl gene ID were converted to human Ensembl gene ID (hg38) with the biomaRt R package (Durinck *et al*, 2005). The search for DNA motif in the promoter regions of deregulated genes was performed using HOMER (Heinz *et al*, 2010) with the findMotifs.pl function and the following parameters: -start -2000 -end 0.

To generate Appendix FigS1B, we used https://rgv.genouest.org/ (Darde *et al*, 2019; Darde *et al*, 2015).

### ChIP-Seq analyses

Snakemake was used for ChIP-Seq analyses. Adaptor were trimmed and filtered using the same parameters than RNA-Seq analyses. Alignment was performed on the mouse genome (mm10) using bowtie2 (Langmead & Salzberg, 2012) with the following arguments: --local. Peak calling was process with MACS2 (Zhang *et al*, 2008) with default parameters against input file to obtains peaks enrichment score of H3K79me2. Mitochondrial peaks and peaks with FDR > 5% were filtered. Peaks were annotated with the R package ChIPseeker (Yu *et al*, 2015) using the annotatePeak() function with the following parameters tssRegion = c(−3000,3000).

### Code availability

The fully reproducible and documented analysis for RNA-Seq and ChIP-Seq is available on github at https://github.com/ManonCoulee/RNAseq_DOT1L_Blanco_2022.

### MS quantification of histone PTMs and of histone variants

To identify histone PTMs, samples were prepared and processed essentially as described in Crespo *et al*. (Crespo *et al*., 2020). Briefly, histones were precipitated by acid extraction from whole testes which were beforehand reduced into powder using a pestle and mortar on dry ice, or from elutriated elongating/condensing spermatids (ES). They were resuspended in loading gel buffer (LDS sample buffer reference 84788, supplemented with NuPAGE sample reducing agent reference NP0009, ThermoFisher) and separated on a 4-12% NuPAGE acrylamide gel (reference NP0321BOX, ThermoFisher). After blue staining (SimplyBlue SafeStain, ThermoFisher), gel bands corresponding to H3 (at about 17 kDa) and H4 (at about 14 kDa) and two gel slices in-between containing H2A and H2B, were cut and then reduced with dithiothreitol, alkylated with iodoacetamide and in-gel digested with 0.1 µg trypsin (V511, Promega) per slice using a Freedom EVO150 robotic platform (Tecan Traging AG, Switzerland). Alternatively, to analyze the relative abundance of histone variants between conditions, histone samples were simply loaded on a gel and migrated for a few mm. The whole zone was cut for robotic in-gel tryptic digestion. The resulting tryptic peptides were analyzed by a liquid chromatography-tandem mass spectrometry coupling made up of a C18 reversed-phase capillary column (75 μm i.d. x 25 cm ReproSil-Pur C18-AQ, 1.9 µm particles, Cluzeau, France) using the UltiMate™ 3000 RSLCnano system (ThermoFisher) coupled to a Q-Exactive HF mass spectrometer (ThermoFisher). The mobile phases consisted of water with 0.1% formic acid (A) and acetonitrile with 0.08% (v/v) formic acid (B). Peptides were eluted with a gradient consisting of an increase of solvent B from 2.8% to 7.5% for 7.5 min, then from 7.5% to 33.2% over 33.5 min and finally from 33.2% to 48% over 6.5 min. Mass spectrometry acquisitions were carried out by alternating one full MS scan with Orbitrap detection acquired over the mass range 300 to 1300 m/z and data-dependent MS/MS spectra on the 10 most abundant precursor ions detected in MS. The peptides were isolated for fragmentation by higher-energy collisional dissociation (HCD) with a collision energy of 27.

Identification of modified peptides was obtained using our in-house established database of histone sequences, MS_histoneDB (El Kennani *et al*, 2017), completed with a list of 500 common contaminants. MS/MS data interpretation was carried out with the program Mascot (http://www.matrixscience.com/) with the following search parameters. The precursor and fragment mass tolerances were 5 ppm and 25 mmu, respectively; enzyme specificity was trypsin; the maximum number of trypsin missed cleavages was set to 5, carbamidomethyl (Cys) was specified as a fixed modification. We indicated as variable PTMs in Mascot acetylation, crotonylation and ubiquitination (more precisely the dipeptide “GlyGly”) of Lys residues, methylation and di-methylation of Lys/Arg and tri-methylation of Lys, as well as N-terminal acetylation of proteins. Filtering of peptide identifications and quantification was performed by using the program Proline (Bouyssie *et al*, 2020). All peptide/spectrum matches of scores below 15 were filtered out; next, all identifications of modified peptides suggested by Mascot for H3 and H4 were visually validated by demanding that a continuous stretch of minimally 5 amino acids be identified in terms of b or y fragment ions, and by ascertaining PTM positioning on Lys/Arg residues. This score threshold is very low, yet it was relevant to use so as to be able to identify very short peptides, such as H3-K18modQLATK. When interpreting our mass spectrometry data against the database MS_histoneDB, we were able to identify and quantify various H3 variants, namely canonical H3 and variants H3.3, testis-specific H3t and H3mm13. H3t differs from canonical H3 at residue 24, while H3.3 and H3mm13 differ from H3 at residues 31 and 29. To estimate the relative abundance of modified histone peptides between samples, normalization to be at constant amount of total H2A, H2B, H3 or H4 was done by dividing the raw MS signals of modified peptides by the sum of the MS signals of modified and unmodified peptides for each histone. To estimate the relative abundance of histone variants between *Dot1l*-KO and CTL samples, variants were quantified by Proline by relying only on peptides specific of these sequences. Samples were normalized to be at constant amount of histone H4. The mass spectrometry proteomics data have been deposited to the ProteomeXchange Consortium via the PRIDE (Perez-Riverol *et al*, 2019) partner repository with the dataset identifier PXD030734 and 10.6019/PXD030734.

### Statistical Analysis

Chi-square and unpaired t-test were used to analyze IVF data. Sperm motility parameters were analyzed using a Mann Whitney tests following angular transformation of percentages. To compare the incidence of sperm head abnormalities, and apoptotic (TUNEL+) spermatids between *Dot1l*-KO and CTL, percentages were converted in angles prior to performing t-tests. Student t-test (corrected for multiple testing when appropriate) was used for all other analyses (fertility, sperm count, testis weight, histology, histone PTM, western blot, and normalized PRM1/2 ratio quantifications). Statistical analyses for RNA-Seq/ChIP-Seq are described in the dedicated paragraphs.

## Data availability

RNA-Seq data have been submitted to ENA repository under project number PRJEB50887 (https://www.ebi.ac.uk/ena/). H3K79me2 ChIP-Seq project number is PRJNA643726. The mass spectrometry proteomics data have been deposited to the ProteomeXchange Consortium via the PRIDE (Perez-Riverol et al., 2019) partner repository with the dataset identifier PXD030734 and 10.6019/PXD030734.

## Acknowledgements

We would like to thank Cochin Institute (INSERM U1016, CNRS UMR8104, Universite Paris Cité) core facilities, in particular, Alain Schmitt from the electron microscopy platform, and all the staff from the animal house, histology (HistIM), genomic (GENOM’IC), cytometry (CYBIO) and imaging (IMAG’IC) core facilities. We would also like to thank Guillaume Meurice for advices on bioinformatics analyses, and Aminata Touré and Marjorie Whitfield for advices and discussion on sperm morphology and motility analyses. M.Cr. and D.P. are grateful to their colleagues in EDyP for their support on LC-MS instruments and in informatics. This work was supported by the Agence Nationale de la Recherche (ANR-17-CE12-0004-01 to J.C., ANR-21-CE44-0035 to D.P.), the Fondation pour la Recherche Médicale (SPF201909009274 to C.G.) and grants from “Ministerio de Economía y competitividad” FI17/00224 to A.I., and “Ministerio de Ciencia e Innovación” PI20/00936 to R.O. and MV20/00026 to A.I. M.B and M.Co. received a PhD funding from Université Paris Cité, M. Cr., from University Grenoble Alps (UGA). The proteomic experiments were partially supported by Agence Nationale de la Recherche under projects ProFI (Proteomics French Infrastructure, ANR-10-INBS-08) and GRAL, a program from the Chemistry Biology Health (CBH) Graduate School of University Grenoble Alpes (ANR-17-EURE-0003).

## Expanded View Figure legends

**Figure EV1.**
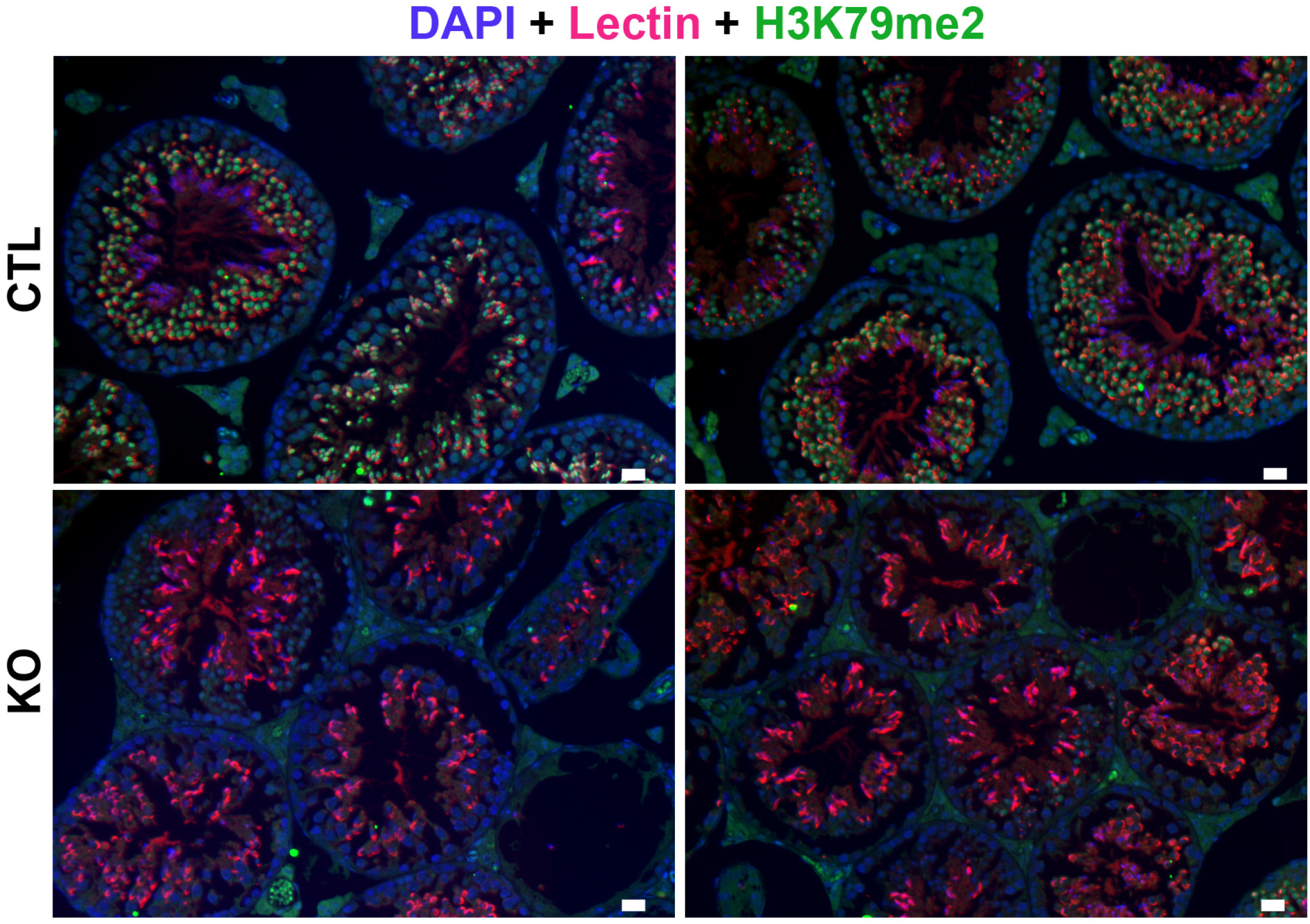
Immunofluorescence detection of H3K79me2 on testicular sections from adult CTL or *Dot1l*-KO mice. H3K79me2 signal is shown in green. DAPI (grey or blue) was used to stain nuclei. Lectin (pink) was used to stain acrosome. Scale bar indicates 20µm.

**Figure EV2.**
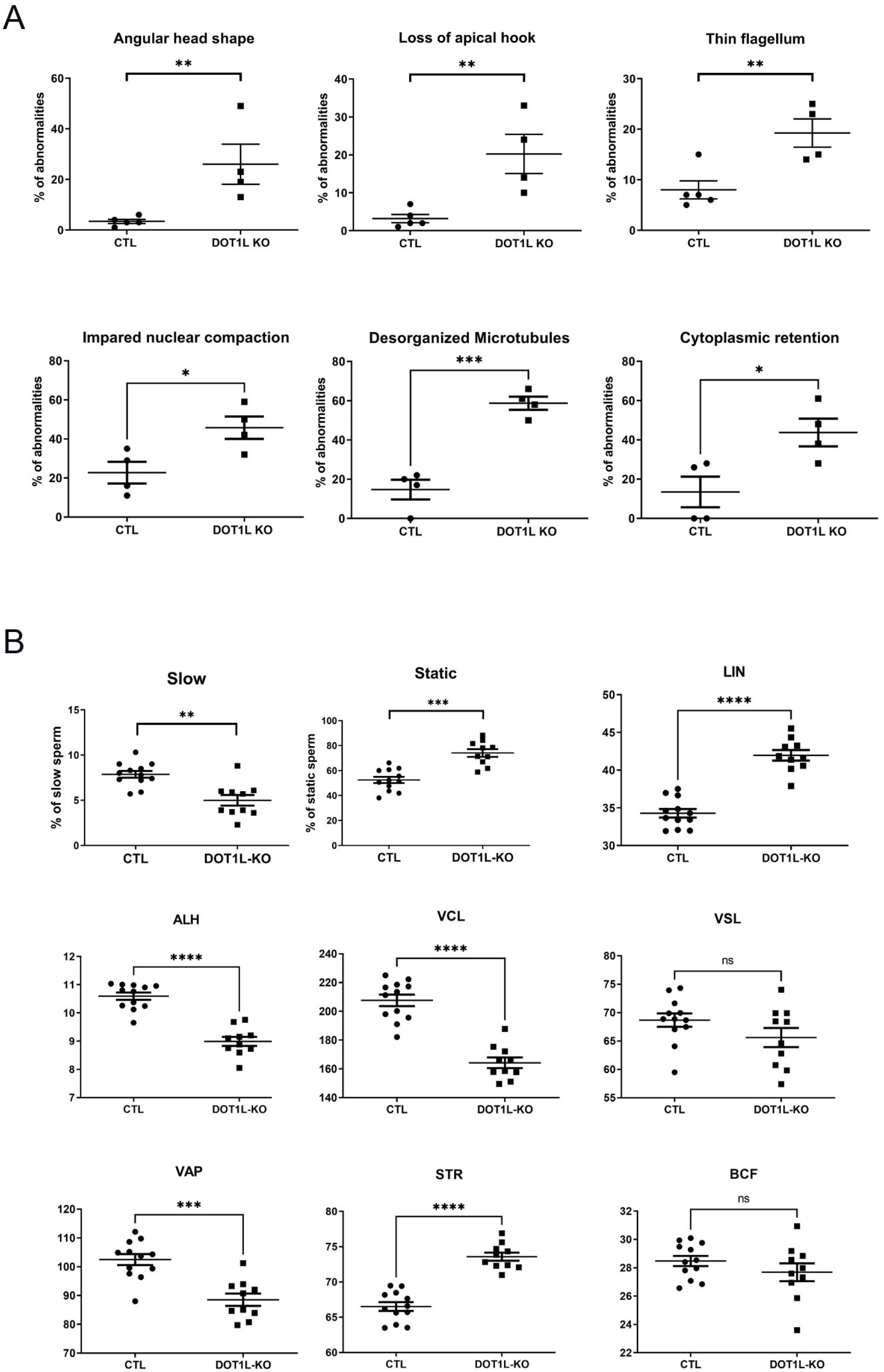
**A) Scatter plots showing the quantification of sperm abnormalities.** Stars indicate significant differences obtained using t-tests after angular transformation of the percentages, with the following p-values. One star indicates a p-value <0.05, two stars a p-value<0.005 and three stars a p-value <0.0005 (N= 4 to 5 CTL and 4 DOT1L-KO males). **B**) **Motility parameters from *Dot1l*-KO and CTL spermatozoa following CASA** (computer-assisted sperm analyses) (N= 12 CTL and 10 KO males). Specifically, *Dot1l*-KO spermatozoa are slower and more static. They have a more linear movement (LIN) and a decreased straightness (STR), decreased lateral displacements of their heads (ALH) with a lower curvilinear velocity (VCL) resulting in a straight-line velocity (VSL) similar to CTL spermatozoa but a higher Average Path Velocity (VAP). Stars indicate significant differences obtained using Mann Whitney t-tests after angular transformation of the percentages (** with a p-value <0.01, *** with a p-value <0.001, **** with a p-value <0.0001, ‘ns’ indicates non-significance). The Beat Cross Frequency (BCF) of CTL and *Dot1l* -KO spermatozoa is similar.

**Figure EV3.**
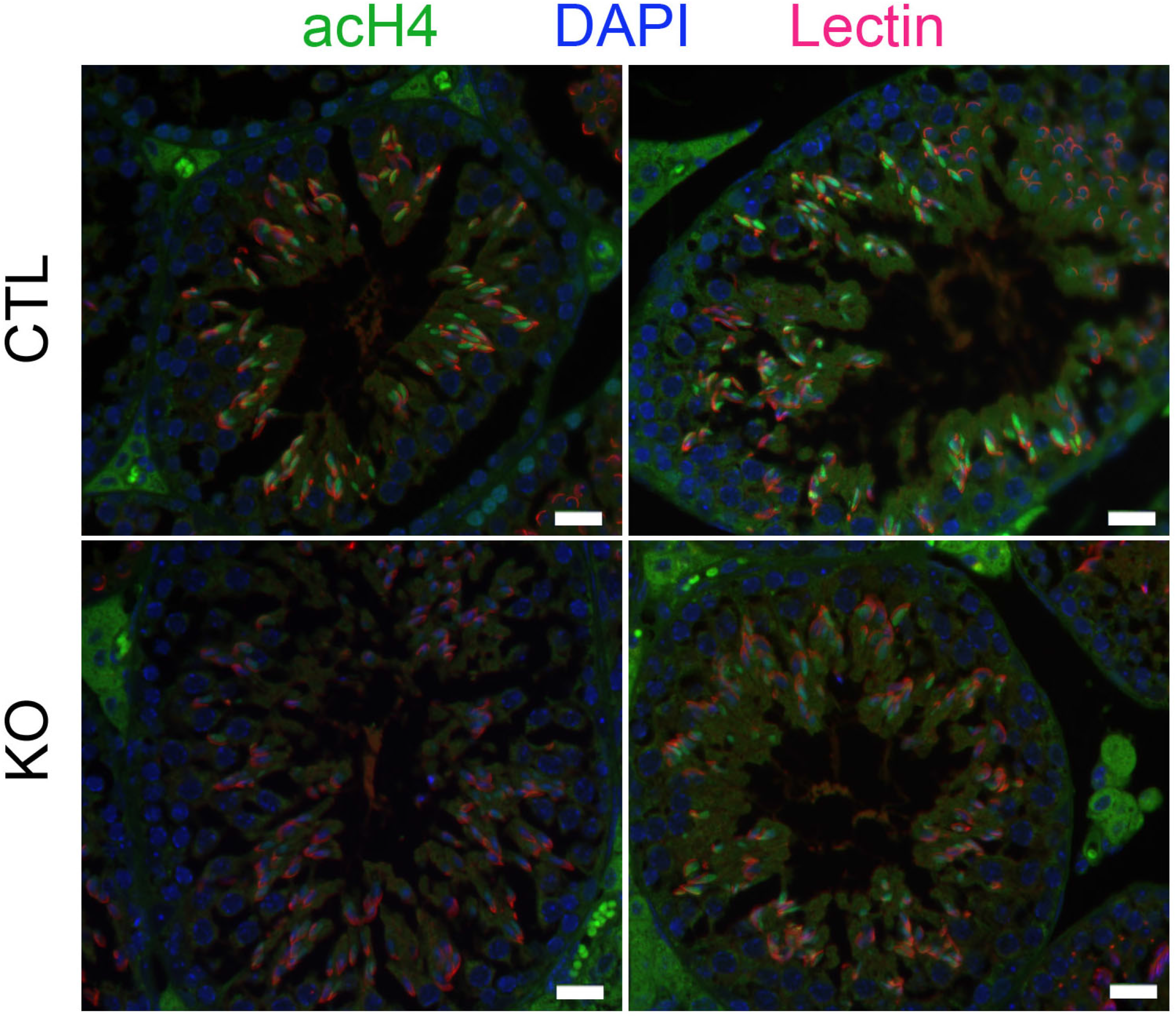
Confirmation of a decrease in H4 acetylation in elongating spermatids (ES) by immunofluorescence. Representative pictures of immunofluorescence detection of acH4 (anti-poly-H4ac, green) on stage XI testicular sections. At this stage, acH4 level is strong in CTL elongating spermatids but appears weaker in *Dot1l*-KO. DAPI (blue) was used to stain nuclei and lectin (red) to stage the acrosome and facilitates tubule staging. Scale bar indicates 20µm.

**Figure EV4.**
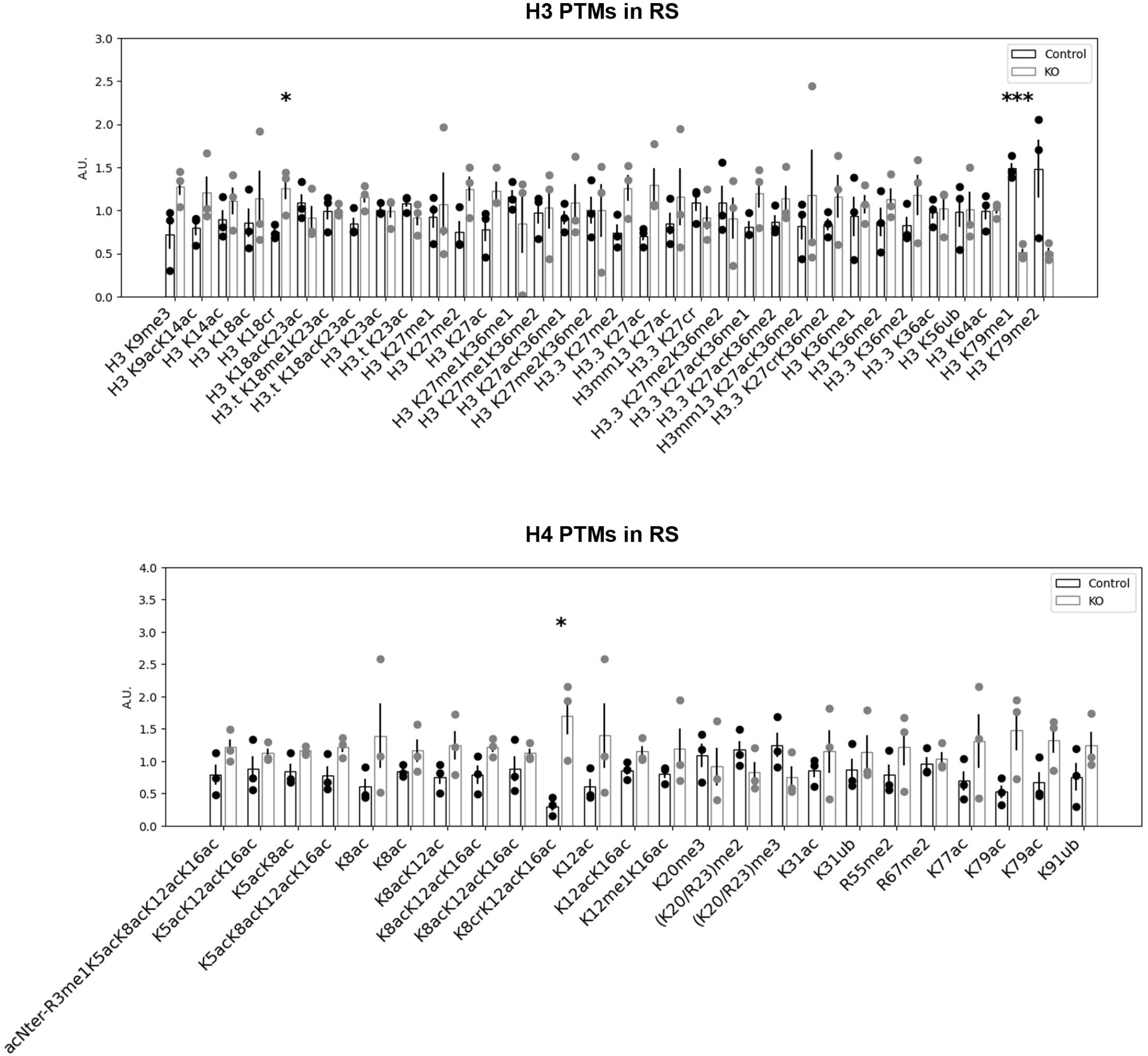
Quantification of histone H3 and H4 PTMs in *Dot1l*-KO and CTL round spermatids (N= CTL and 3 KO). LC-MS/MS data acquired on H3 and H4 were processed as described in Figure 4. One star indicates a p-value <0.05, two stars a p-value<0.005 and three stars a p-value <0.0005, obtained with t-tests.

**Figure EV5.**
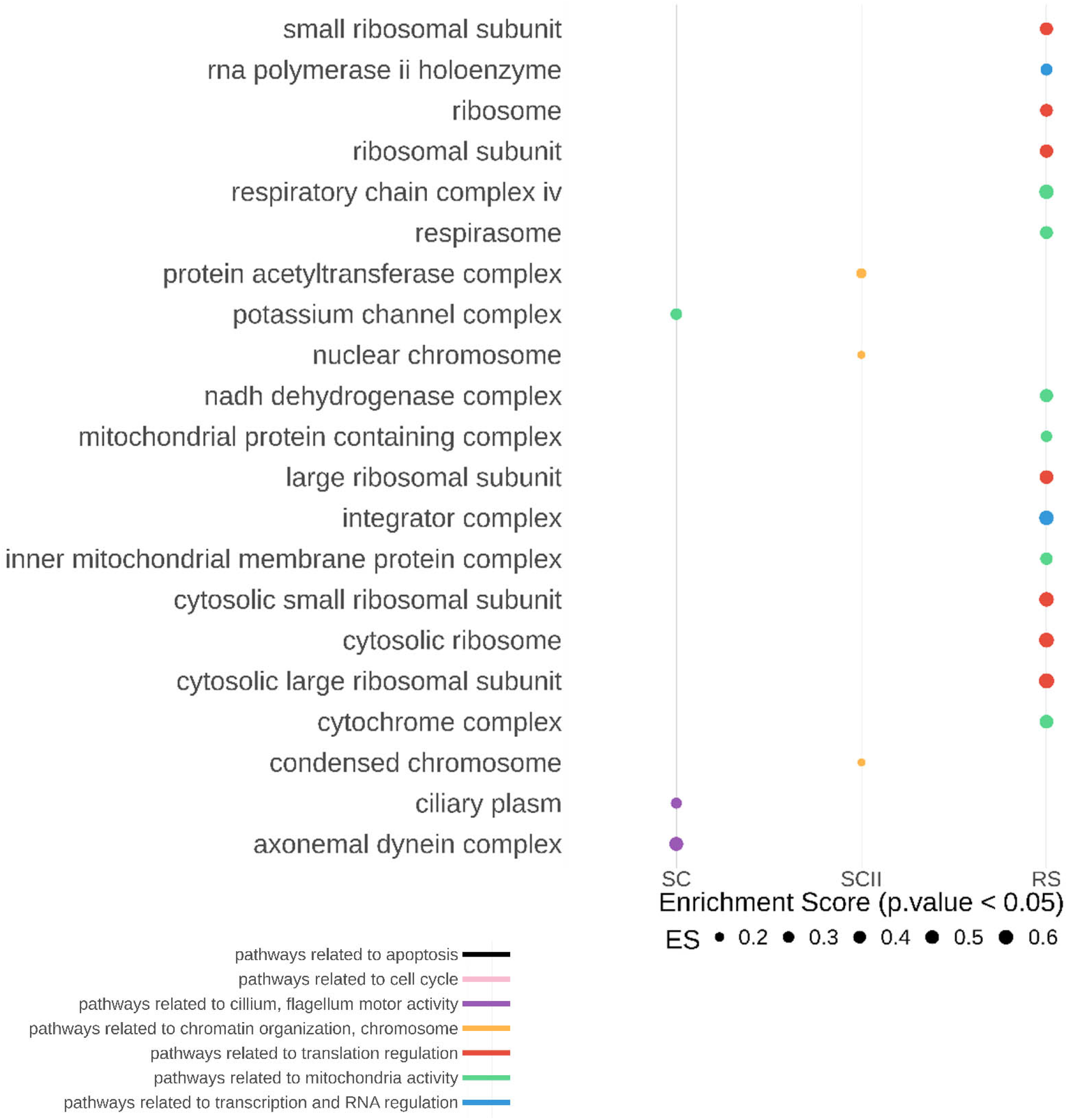
GSEA analysis of RNA-seq data from *Dot1l*-KO vs CTL SC, SCII and RS. The figure shows all the “cellular components” found significantly downregulated in *Dot1l*-KO primary spermatocytes (SC), secondary spermatocytes (SCII) and round spermatids (p<0.05), ranked by their enrichment score (ES).

