## Appendix figures and tables for "DOT1L regulates chromatin reorganization and gene expression during sperm differentiation"

**Appendix Figure 1. Complement to Figure 1. A) RNA expression overview of *DOT1L* RNA in human and mouse tissues**. Data were obtained from The Human Protein Atlas [https://www.proteinatlas.org/; consensus dataset, consisting of normalized expression (nTPM) levels for 55 tissue types, created by combining the HPA and GTEx transcriptomics datasets] or from mouse tissues from mouse ENCODE transcriptome data (Data ref: Mouse ENCODE transcriptome data, NCBI Sequence Read Archive PRJNA66167, 2011). **B) Expression of *Dot1l* mRNA based on single cell RNA-seq analyses from adult testicular cells** (Data ref: (Green *et al*, 2018) analyzed using https://rgv.genouest.org/ (Darde *et al*, 2019; Darde *et al*, 2015)). **C) Summary of DOT1L protein expression in mouse germ cells based on immunofluorescence detection on testicular sections. Scheme modified after Russell et al. (Russell *et al*, 1990).** DOT1L protein is detected in all germ cells but is particularly abundant in late spermatocytes and in round spermatids. **D) Generation of *Dot1l* KO allele.** Drawing representing the beginning of mouse *Dot1l* gene (from exon 1 to exon 3), the location of loxP sites and genotyping primers. Bottom right insert shows typical pictures of products from PCR using *Dot1l* genotyping primers (F+R1+R2) following run on a 2% agarose gel. **E) Estimation of *Dot1l* knockout efficiency by western blot (uncropped western blot image of Figure 1B).** Detection of DOT1L and TUBULIN in whole testicular protein extracts from CTL and *Dot1l*-KO mice. The most abundant and high molecular weight bands are DOT1L testis-specific (~185KDa) and canonical proteoform. Additional bands most likely corresponding to smaller, less abundant proteoforms (such as Q679P5 at ~122KDa or Q6XZL7 at ~68KDa) are also visible in case of long exposure. All DOT1L proteoforms are markedly reduced in *Dot1l*-KO testicular extracts. On the right is shown a graphic representation of the quantification of DOT1L signal (all bands) normalized to TUBULIN signal**. F) Immunohistochemistry detection of DOT1L in testicular sections from CTL and *Dot1l*-KO (KO) adult mice counterstained with hematoxylin (complement panel to Fig 1C).** SC = Spermatocytes, RS = Round spermatids, ES = Elongating spermatids. Pictures were taken using the same parameters. Scale bars indicate 50 µm. **G) Estimation of *Dot1l* knockout efficiency by immunofluorescence.** Representative pictures of immunofluorescence detection of DOT1L (Abcam 64077 antibody, in green) on testicular sections from adult CTL or KO mice. DAPI (grey or blue) was used to stain nuclei; Lectin (pink) was used to stain acrosome. Scale bar indicates 20µm. In CTL testicular sections, tubules containing round spermatids are uniformly stained (bottom panel). In *Dot1l*-KO sections, the majority of tubules do not show any DOTL1 protein signal (top panel) but in some tubules, DOT1L signal can still be visible (middle panel). Below the immunofluorescence pictures is shown a graphic representation of the percentage of round spermatids with a visible DOT1L signal. N=5 CTL, 3 HET and 6 KO males between 2.5-4 month old (6 to 10 tubules per individual). Quantification was performed on stages I to V tubules. **H) Verification of *Stra8-Cre* recombinase activity and specificity in adult mouse testis.** The Cre reporter mouse line *mTmG* carries a *GFP* transgene expressed upon Cre induction. In this transgene, GFP is targeted to the cell membrane (Muzumdar *et al*, 2007). In testicular sections from *mTmG* mice (left panel), using an anti-GFP antibody, no signal is detected, excepted for faint auto-fluorescence from Leydig cells (asterisk). In *mTmG ; Stra8-Cre* mice (right panel) the Cre recombinase is specifically expressed in germ cells and GFP (in green) is detected in most germ cells. DAPI (grey) has been used to stain cell nuclei. Top right panel represents a 2X magnification. **I) Uncropped western blot image of Fig 1D.** Note that the membrane was cut at ~70KDa and ~35KDa to generate three pieces which were incubated with anti-DOT1L antibody (top part), with anti-TUBULIN antibody (mid part) or with anti-H3K79me2 antibody (bottom part).

**Appendix Figure 2. Complement to Figure 2. Detailed analyses of reproductive parameters from *Dot1l*-KO males and CTL**. 'cah' indicates animals produced in a conventional animal house. 'spf' indicates animals produced in the specific pathogen-free animal house. HET are heterozygous animals of the following genotype *Dot1l*$^{Fl/\Delta}$. Obtained p-values (following unpaired t-tests) are indicated above; "ns" indicates non-significant difference. **A**). Average testis weight (normalized to body weight, mean value + standard error of the mean) in different groups of males (N=9 for CTLcah, N=6 for HETcah, N= 9 for DOT1Lcah, N=12 for CTLspf, N=2 for HETspf and N= 17 for DOT1Lspf). **B**) Average number of epididymal spermatozoa (mean value + standard error of the mean) in different groups of males (N=12 for CTLcah, N=6 for HETcah, N=12 for DOT1Lcah, N=9 for CTLspf, N=2 for HETspf and N= 13 for DOT1Lspf). **C**) Graphic representation of the percentage of abnormal tubules in *Dot1l*-KO (N=5) CTL (N=6) and HET (N=4) testes (from both cah and spf). Obtained p-values (following unpaired t-tests on empty tubules values) are indicated above. **D**) and **E**) Results from the tests of fertility (natural mating in spf with wild type females for ~3 months) of *Dot1l*-KO, HET and CTL males (N=7 for KO, N= 2 for HET and N=6 for CTL). **F**) Schematic diagram representing the frequency of each cell type in CTL and *Dot1l*-KO testes, as calculated by FACS. 4N = Primary spermatocytes, 2N = Secondary spermatocytes, N = spermatids, SP = side population representing premeiotic germ cells (i.e. spermatogonia). **G)** Results from *in vitro* fertilization assays. The percentage of fertilized oocytes is indicated for each male. Values obtained for *Dot1l*-KO spermatozoa are significantly different from WT or CTL spermatozoa (chi-square, p<0.01).

**Appendix Figure 3. Complement to Figure 3. A) Sperm vitality**. Bar graph showing sperm vitality measured by eosin-nigrosine staining in which dead spermatozoa appear violet. Sperm samples from 4 CTL and 4 *Dot1l*-KO males were analyzed. **B) *Dot1l*-KO sperm chromatin is sensitive to nucleoplasmin-induced decompaction.** Quantification of histone H3 levels following NPM (nucleoplasmin) partial decondensation of CTL and *Dot1l*-KO (KO) epididymal spermatozoa. Quantification was performed on the two fractions obtained after centrifugation: (i) the supernatant which contains histones from the NPM-de-compacted chromatin, and the pellet which contains compact chromatin, i.e. "resistant" to the NPM treatment we used. Left panel: Bar graph showing quantification results of supernatant/pellet levels in 3 independent experiments (E1, E2 and E3, which are technical and biological replicates). Right panel: Representative image of western blot using anti-histone H3 antibody (PanH3). PAnH3 is directed against the C-terminal part histone H3 and detects full length histone H3 at 17 kDa (black arrow, H3-C) and its two cleaved forms previously described (Yamaguchi et al., 2018): one occurring between Ala21-Thr22 (red arrow) and the other one between Thr22-Arg26 (the smallest form at 13 kDa, green arrow). **C)** Uncropped western blot images from TH2B and H3 detections in spermatozoa (shown in Fig 3E), and corresponding Ponceau staining of the membranes. A non-specific band is visible in all lanes with Ponceau; it indicates that similar amounts of material were loaded in all lanes. **D)** Comparison of TH2B and H3 levels in the same CTL (black) and *Dot1l*-KO (blue) samples. The graph shows a good correlation between TH2B and H3 quantifications. **E)** Top panel: uncropped western blot images from TNP2 detection in spermatozoa (shown in Fig 3F), and corresponding Ponceau staining of the membranes. A non-specific band is visible in all lanes with Ponceau; it indicates that similar quantity of material was loaded in all lanes. Bottom panel: immunofluorescence detection of TNP2 (green) on stage VI-VIII testicular sections. At this stage, the chromatin of condensed spermatids is packaged with protamines, and TNP2 signal is not (or weak) in CTL. In *Dot1l*-KO, some condensed spermatids (inside of the tubule) have a stronger signal. DAPI (blue) was used to stain nuclei. **F)** Box plot of the raw PRM1/PRM2 ratio following acid urea gel electrophoresis (mean + standard error of the mean) complement to Fig 3G. A.U. = arbitrary units. One star indicates a p-value <0.05 (unpaired t-test).

**Appendix Figure 4. Complement to Figure 4. Quantification of histone PTMs and variants in *Dot1l*-KO and CTL samples.** One star indicates a p-value <0.05, two stars a p-value<0.005 and three stars a p-value <0.0005, obtained with t-tests. **A) Mass spectrometry quantification of H3 and H4 PTMs in whole testes (N= 4 CTL and 4 KO).** After normalization to be at constant amount of histone H3 or H4 in each analyzed sample (see Material and methods), mass spectrometry signals were divided by the average signal in both conditions (CTL and *Dot1l*-KO), so as to be able to represent all peptides in the same figure, whatever their MS intensity. Error bars represent standard deviations calculated on the measurements made on biological replicates. Each modified peptide identified and quantified by MS analysis is defined as a list of PTMs separated by commas. Some modified sites were detected and quantified in several tryptic peptides due to possible missed cleavages (e.g. H4K20me1 was quantified in K20VLRDNIQGITK, K20VLRDNIQGITKPAIR and K20VLRDNIQGITKPAIRR); we indicated the measurements made on such peptides with the same label (K20me1 in the aforementioned example), which allows verifying the coherence of quantitative measurements. In a few instances, a precise localization of the modification site was not possible from the MS/MS spectra. This was typically the case between H4R19, K20 and R23 on which a methyl (me1) and dimethyl (me2) were detected. We indicated such uncertainties by (R19/K20/R23)me1 and (R19/K20/R23)me2, respectively. **B) Mass spectrometry quantification of H2A and H2B in elongating/condensing spermatids (ES) (N=3 CTL and 3 KO).** LC-MS/MS data acquired on H2A and H2B were processed as described above for H3 and H4 in whole testes, except that MS signals detected on modified peptides were normalized to be at constant amount of H2A or H2B in each analyzed sample. **C) Mass spectrometry quantification of histone variants in elongating/condensing spermatids (ES) and in round spermatids (RS).** The abundance of each histone variant was estimated from the sum of intensities of its identified specific proteolytic peptides. The abundance of each protein was then normalized to be at constant amount of histone H4 between samples. For some variants, MS values were divided by 10, so as to be able to visualize all the proteins on the same figure, whatever their MS intensity. Error bars represent standard deviations calculated on the measurements made on biological replicates. **D) Confirmation of a decrease in H4 acetylation in elongating spermatids (ES)**. Western blot detection of H4K16ac, poly-H4ac and H3. Higher molecular weight bands likely represent non-denatured forms of histone H4. Western blot normalization was performed by detecting the same membranes with anti-H3 antibody. Anti-H3 rather than anti-H4 antibody was chosen to avoid interference with acetylated H4 signal. Histones H3 and H4 are organized as obligate tetramers (or occasionally heterodimers) in nucleosomes; therefore, they are expected to be present in the chromatin in the same quantity. **E) Mass spectrometry quantification of histone H2A and H2B PTMs in round spermatids (N= CTL and 3 KO).** LC-MS/MS data acquired on H2A and H2B were processed as described above for ES. **F) Mass spectrometry quantification of histone H3 PTMs in spermatozoa (N= 4 CTL and 4 KO).** Same legend as above. H4 PTMs levels were too weak to be quantified.

**Appendix Figure 5. Complement to Fig 5. A)** Upset plot showing the number of deregulated genes in *Dot1l*-KO vs CTL primary spermatocytes (SC), secondary spermatocytes (SCII) and round spermatids (RS) obtained using DEseq2 differential gene expression analyses. Top panel: with Fold Change > 1.5 and FDR < 0.05. Bottom panel: with Fold Change>1.5 and p value <0.05 (FDR was calculated *a posteriori* on the selected gene lists and found to be <0.05). **B)** Bar graph showing the percentage of deregulated genes in *Dot1l*-KO RS found with the two types of differential analyses and which are enriched in H3K79me2 at their gene body in WT RS. **C)** List of *Slc* genes found deregulated in *Dot1l*-KO round spermatids. **D)** Top: results from HOMER motif analysis on the promoter (2kb) of upregulated genes (p-value < 0.05) in KO RS (top 10). BCL6/BCL6B binding DNA motif is the most significant of the identified motifs (and the only one which does not appear as “possible false positive”). Bottom: list of genes identified with HOMER which have BCL6 DNA binding site in their promoter.

### Appendix Figure S1

## S1A

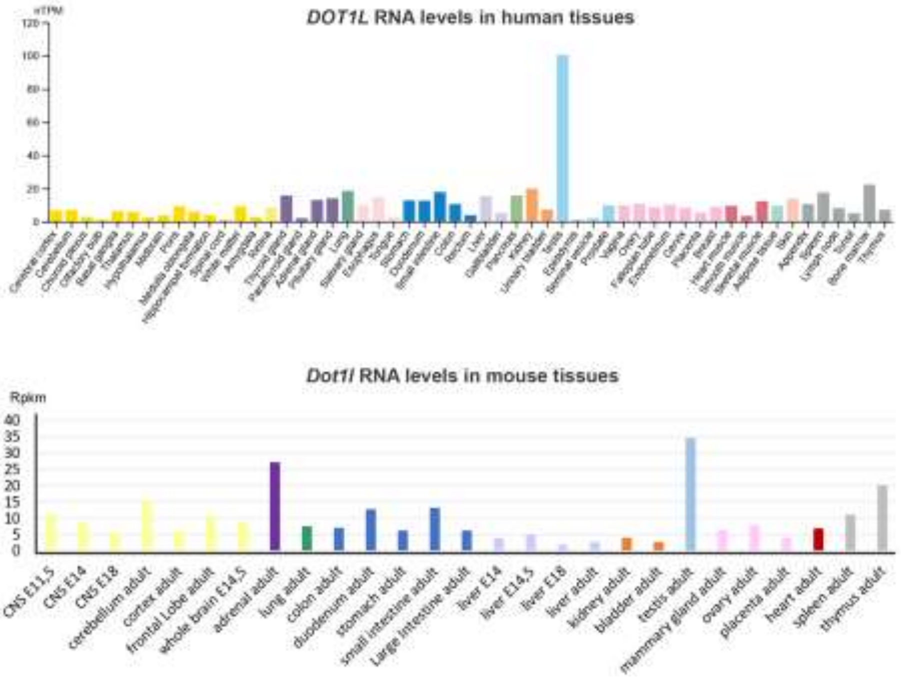

## S1B

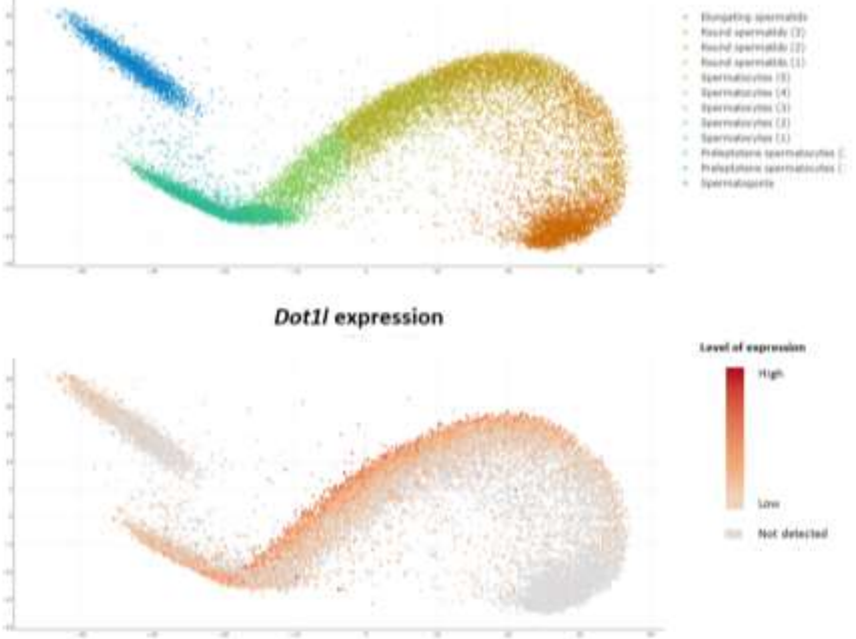

## S1C

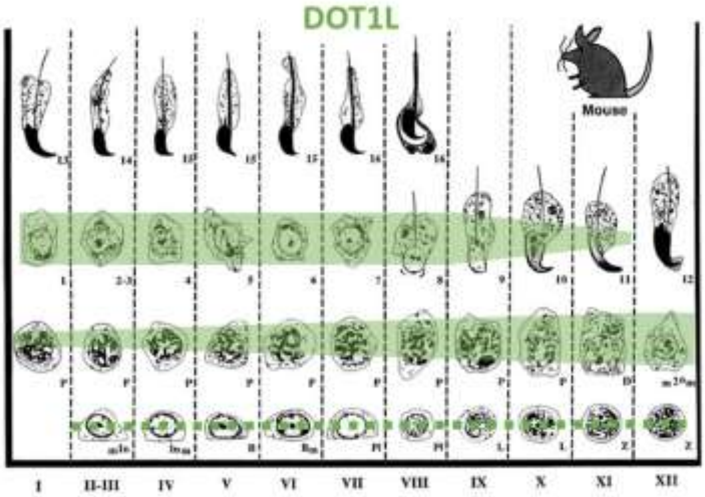
DOT1L
Mouse
I
II-III
IV
V
VI
VII
VIII
IX
X
XI
XII

## S1D

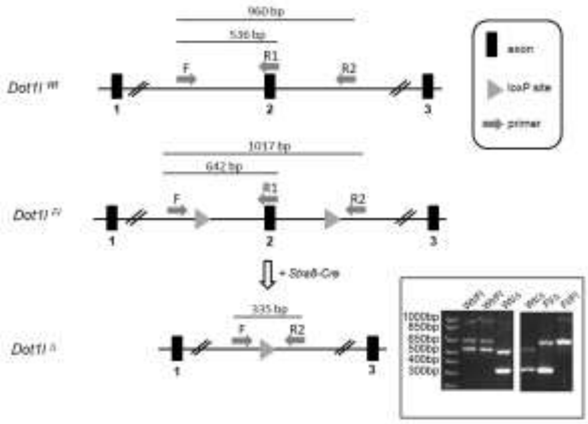
960 bp
536 bp
F
R1
R2
exon
loxP site
primer
1017 bp
642 bp
335 bp

## S1E

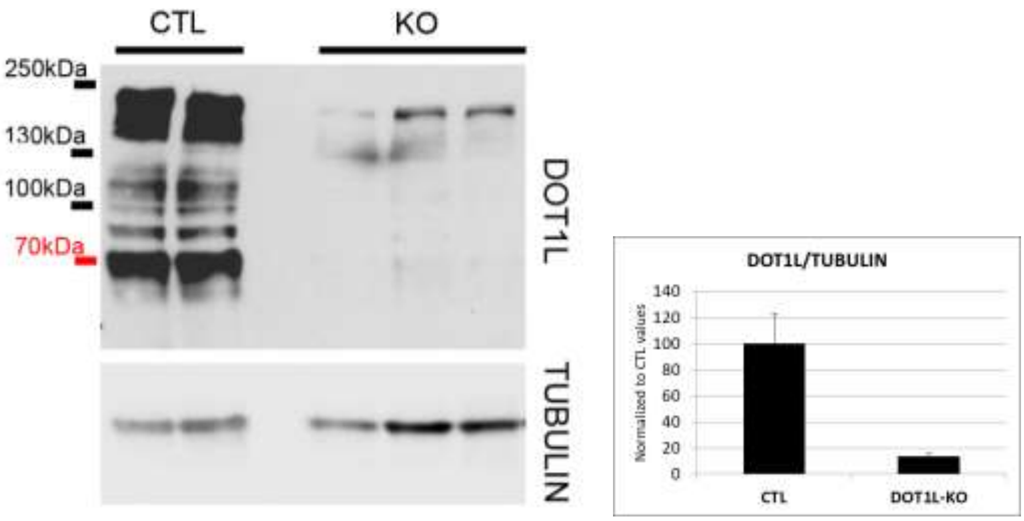
CTL
KO
250kDa
130kDa
100kDa
70kDa
DOT1L
TUBULIN
DOT1L/TUBULIN
Normalized to CTL values
140
120
100
80
60
40
20
0
CTL
DOT1L-KO

## S1F

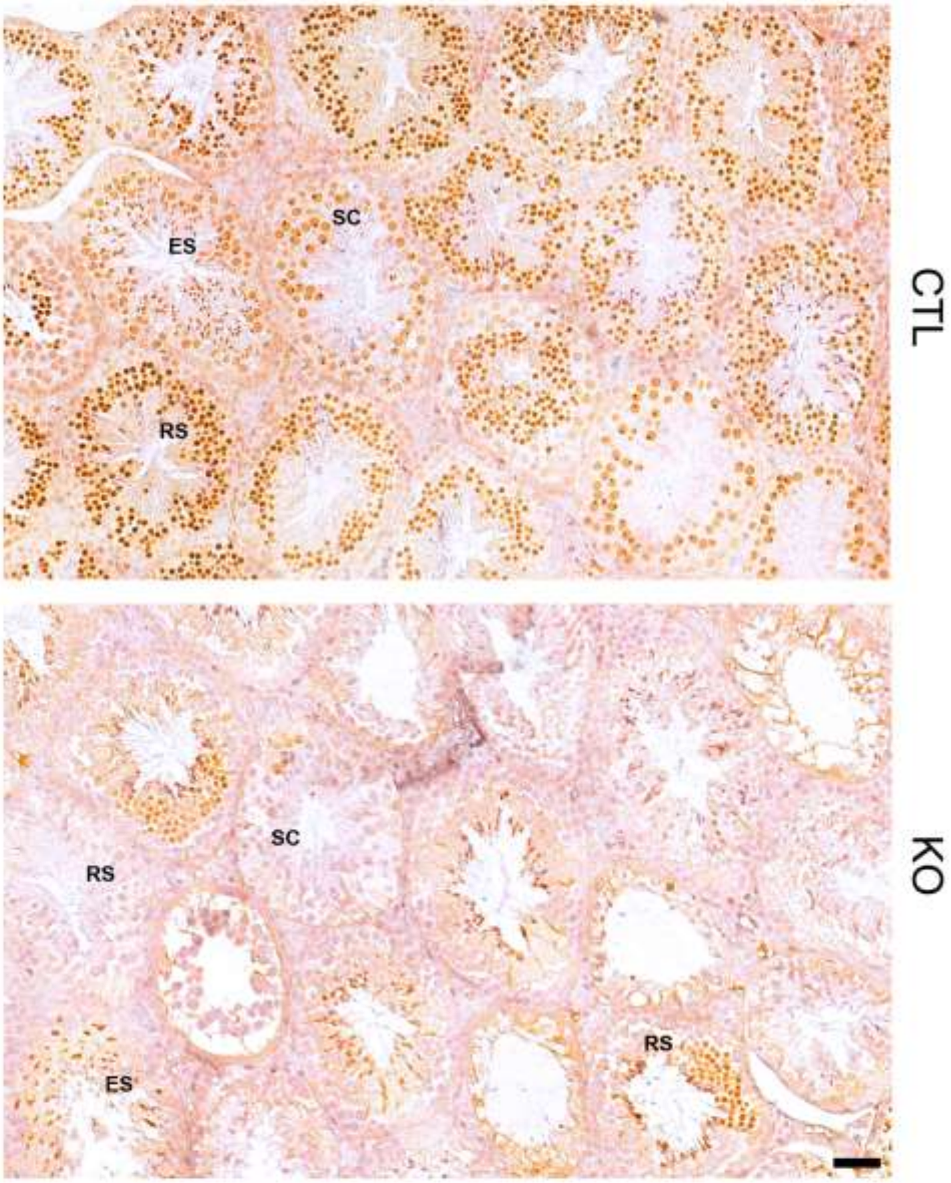

## S1G

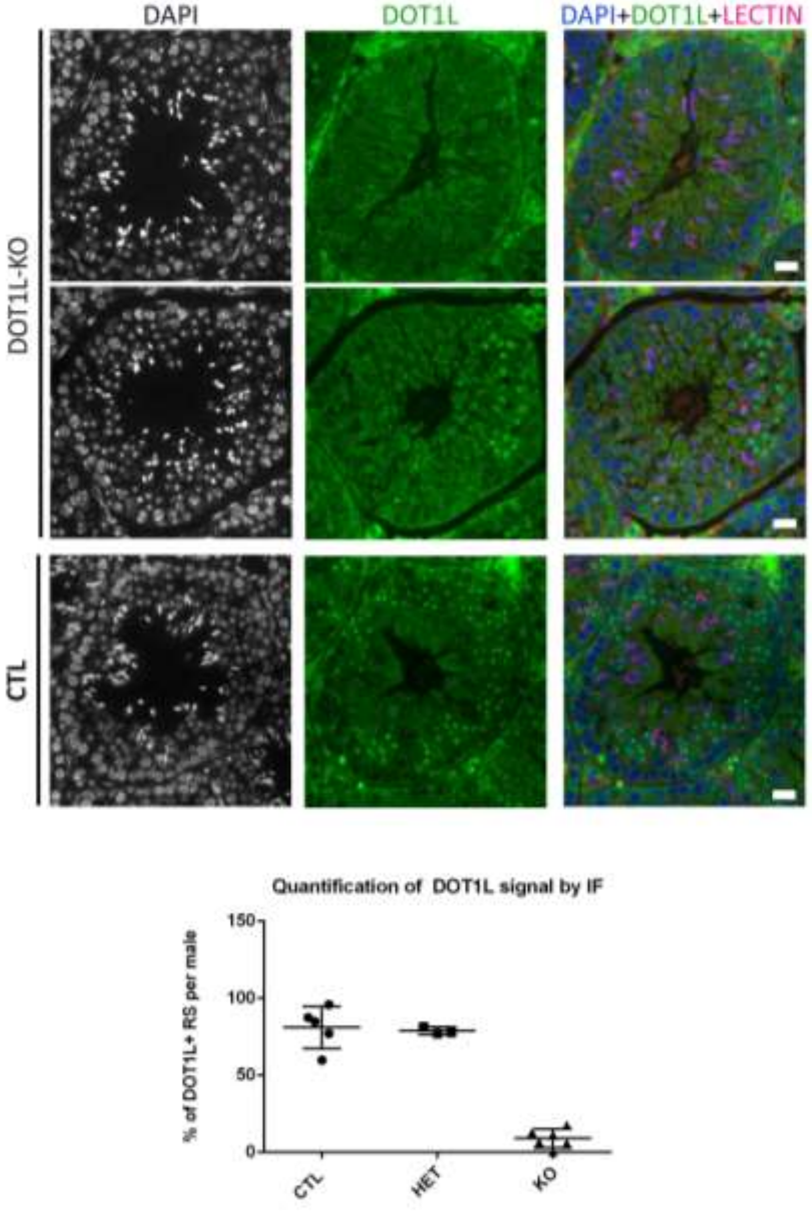
DAPI
DOT1L
DAPI+DOT1L+LECTIN
DOT1L-KO
CTL
Quantification of DOT1L signal by IF
% of DOT1L+ RS per male
150
100
50
0
CTL
HET
KO

## S1H

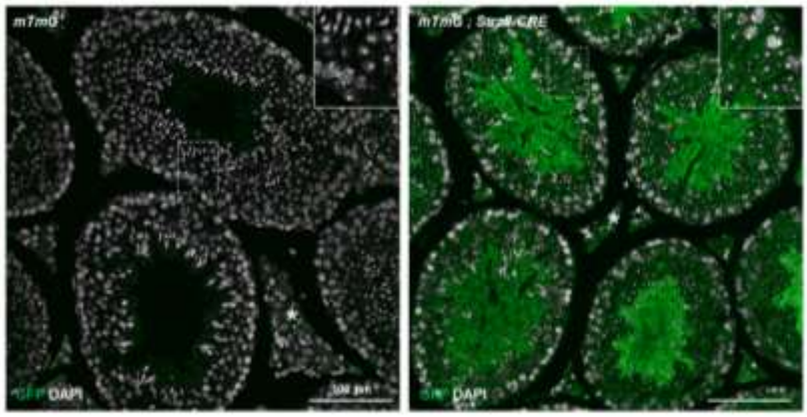
DAPI
DAPI

**S1I**

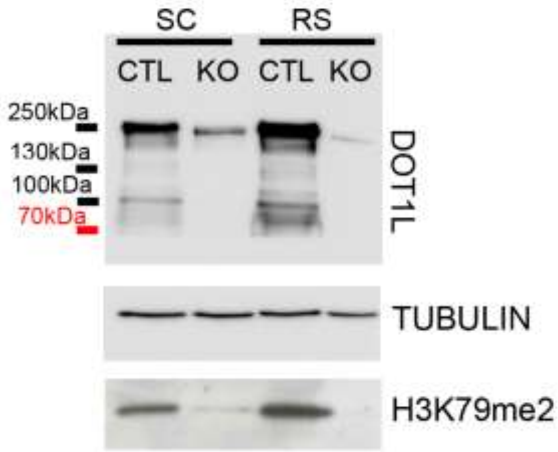
SC
RS
CTL
KO
CTL
KO
250kDa
130kDa
100kDa
70kDa
DOT1L
TUBULIN
H3K79me2

### Appendix Figure S2

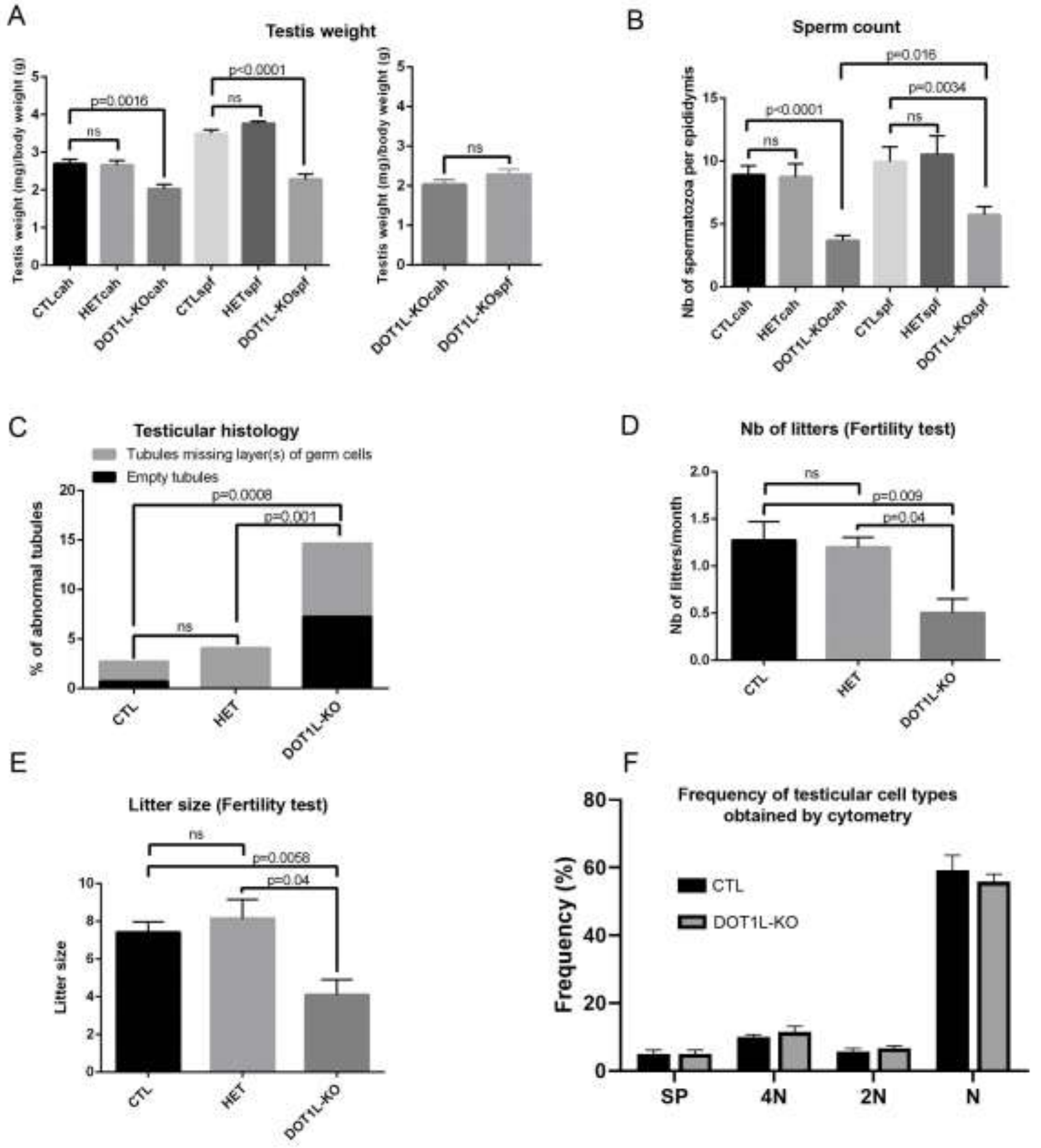

G

**In vitro fertilization**

| | % of oocytes in vitro fertilized (nb of fertilized/total) |
|---|---|
| WT 1 | 52.4 (11/21) |
| WT 2 | 34.8 (24/69) |
| CTL 1 | 36.7 (18/49) |
| CTL 2 | 21.9 (14/64) |
| HET 1 | 37.5 (9/24) |
| DOT1L-KO 1 | 0 (0/37) |
| DOT1L-KO 2 | 0 (0/29) |
| DOT1L-KO 3 | 0 (0/27) |
| DOT1L-KO 4 | 0 (0/87) |

### Appendix Figure S3

## S3A

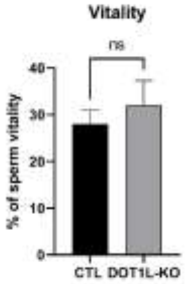

## S3B

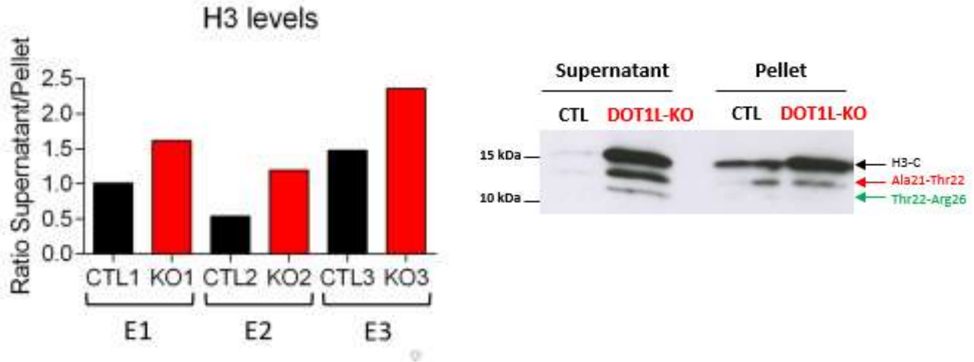

## S3C

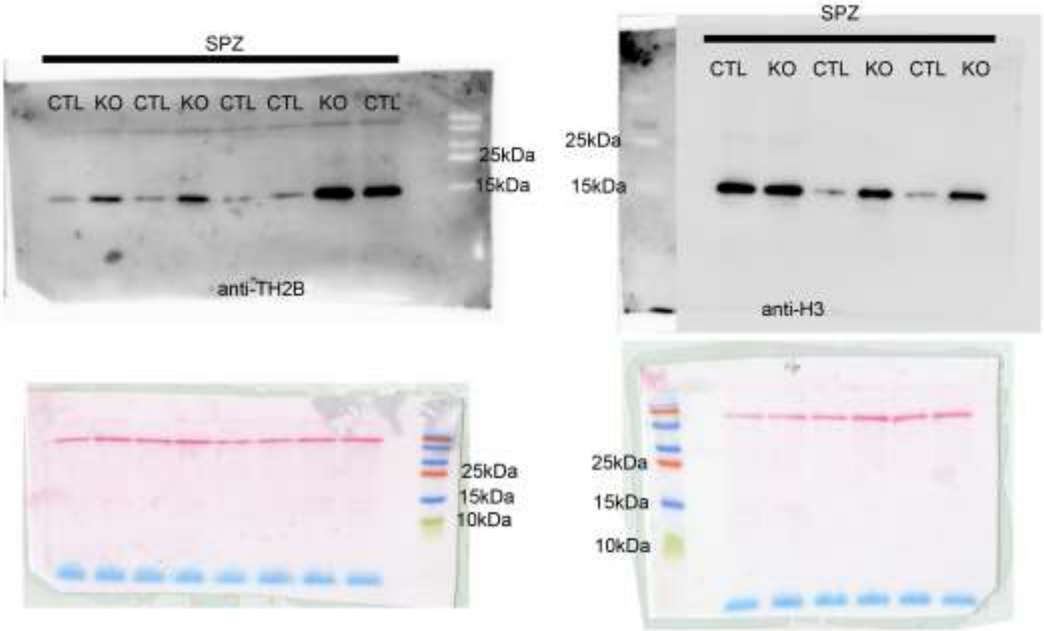

## S3D

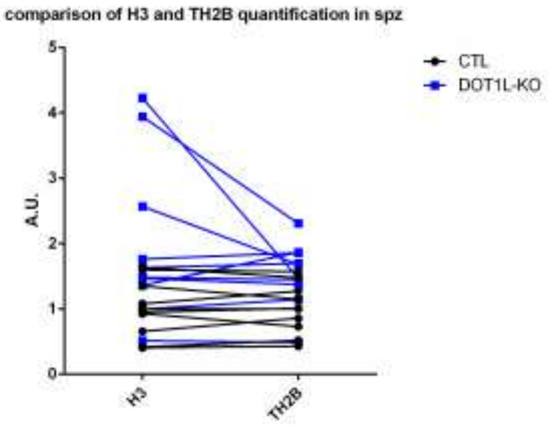

**S3E**

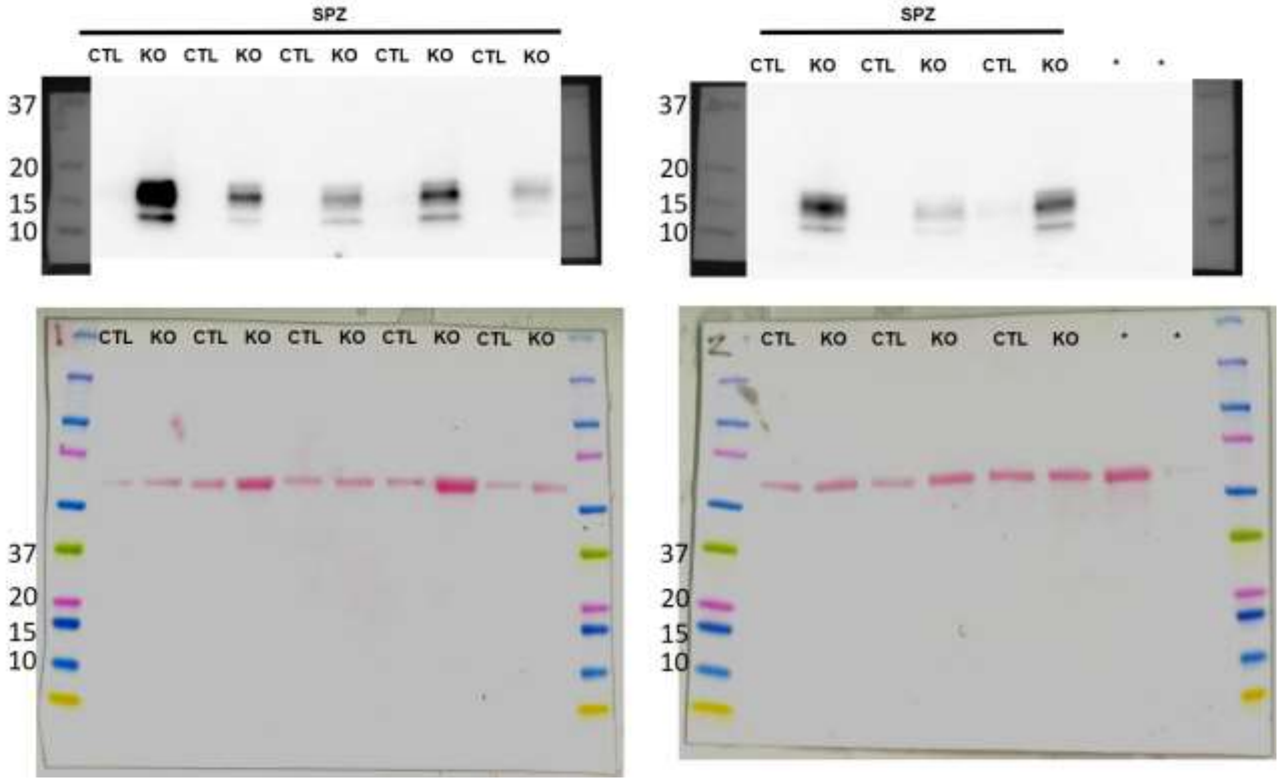
SPZ
CTL KO CTL KO CTL KO CTL KO CTL KO
37
20
15
10
SPZ
CTL KO CTL KO CTL KO * *
37
20
15
10
CTL KO CTL KO CTL KO CTL KO CTL KO
37
20
15
10
CTL KO CTL KO CTL KO * *
37
20
15
10

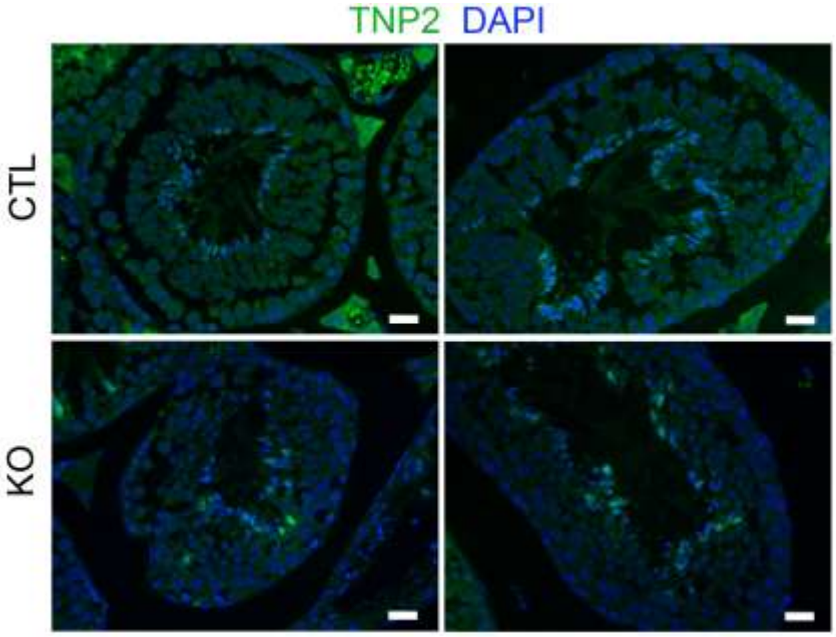
TNP2 DAPI
CTL
KO

# S3F

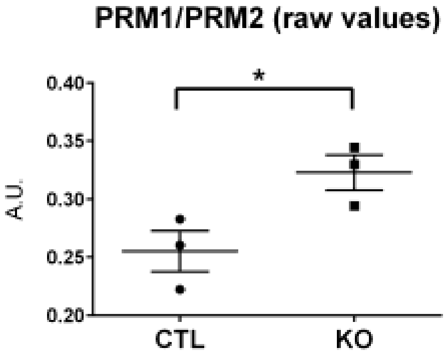

### Appendix Figure S4

## S4A

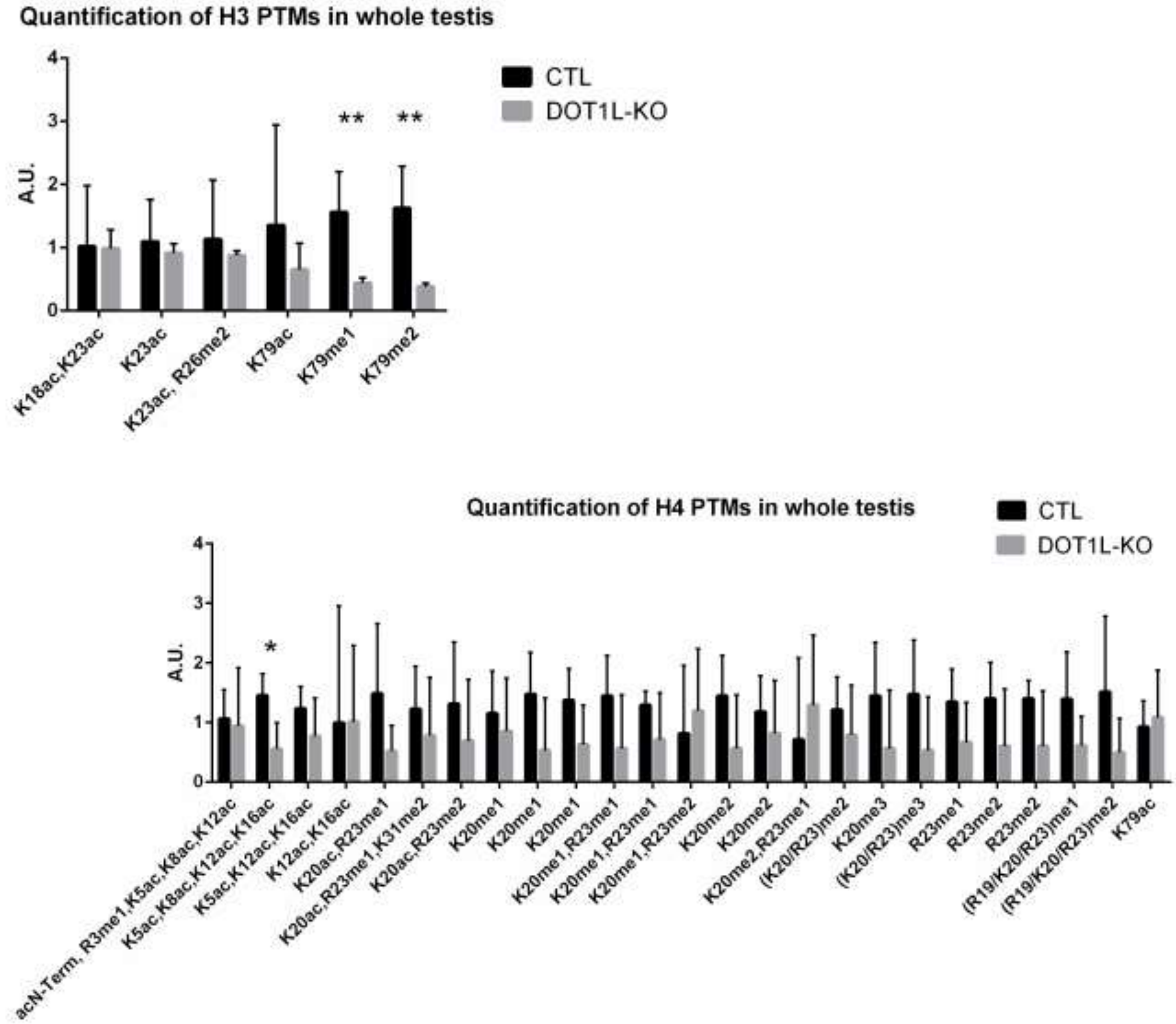

## S4B

**H2A PTMs in ES**

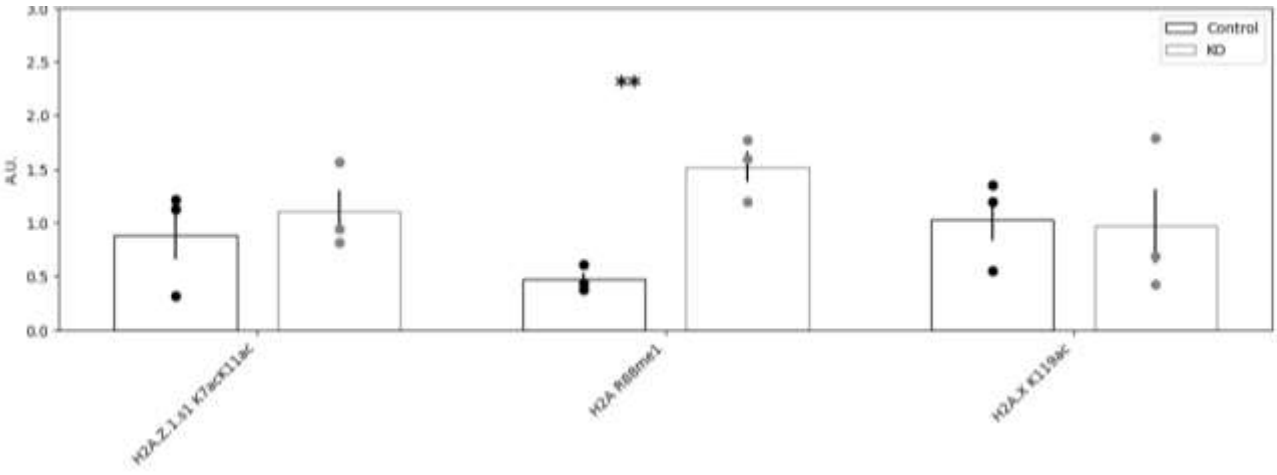

**H2B PTMs in ES**

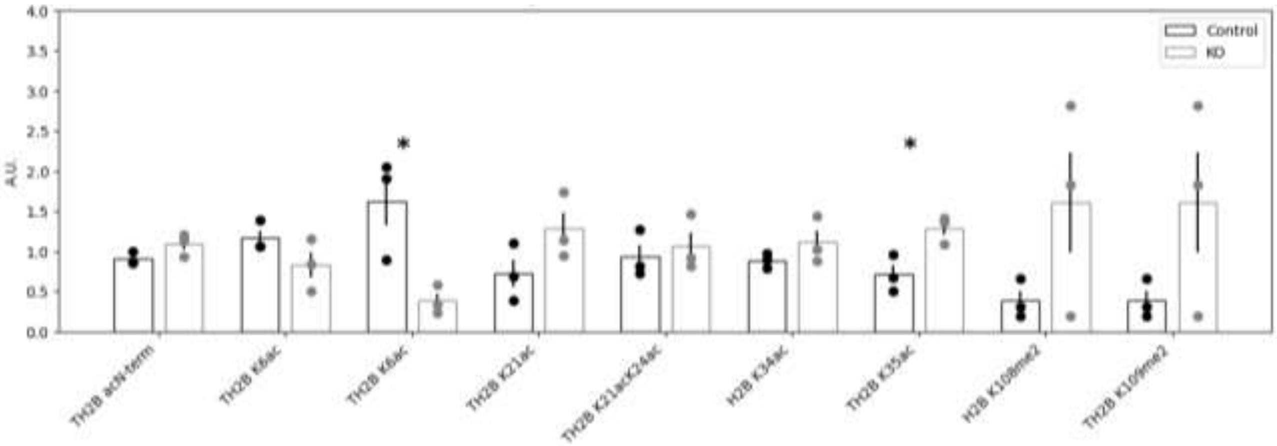

## S4C

**Histone variant levels in ES**

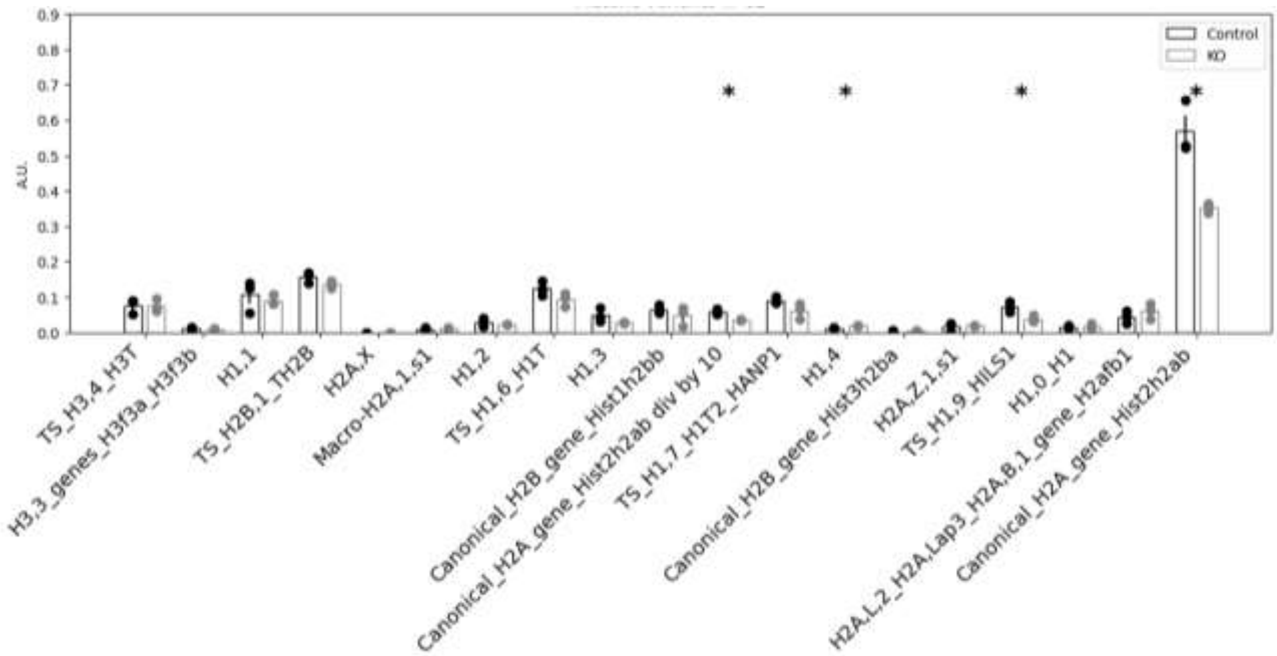

**Histone variant levels in RS**

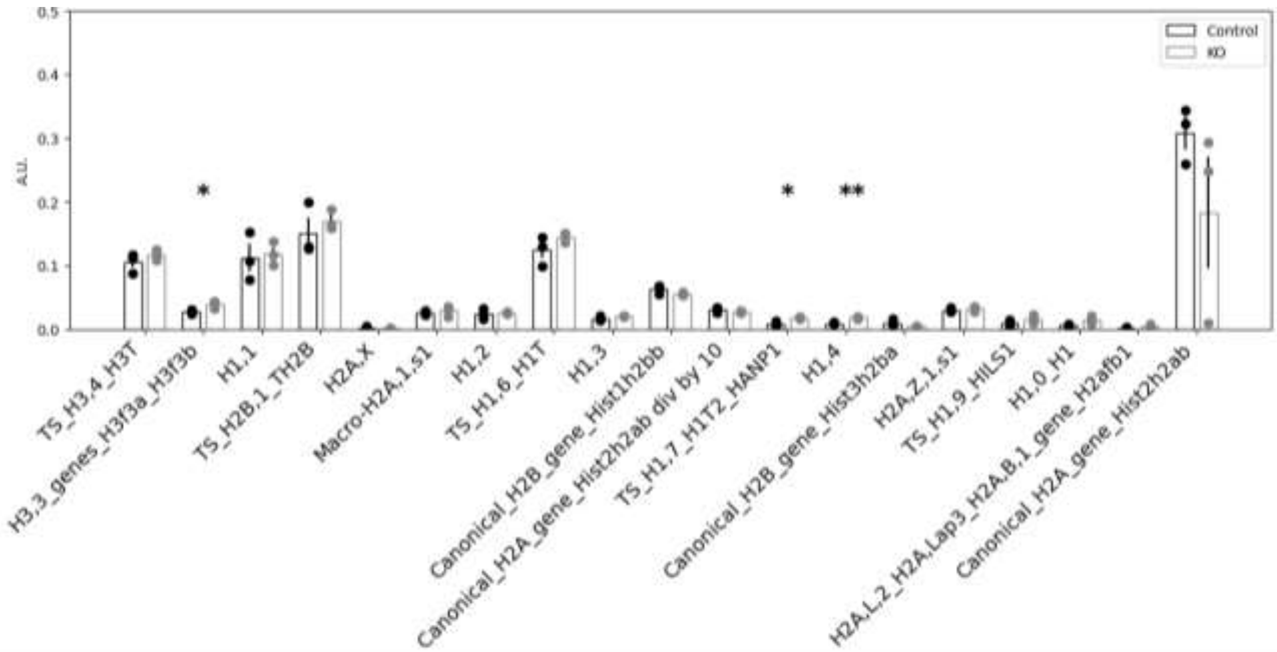

## S4D

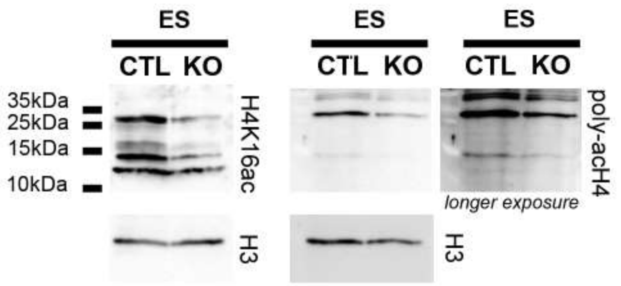

## S4E

**H2A PTMs in RS**

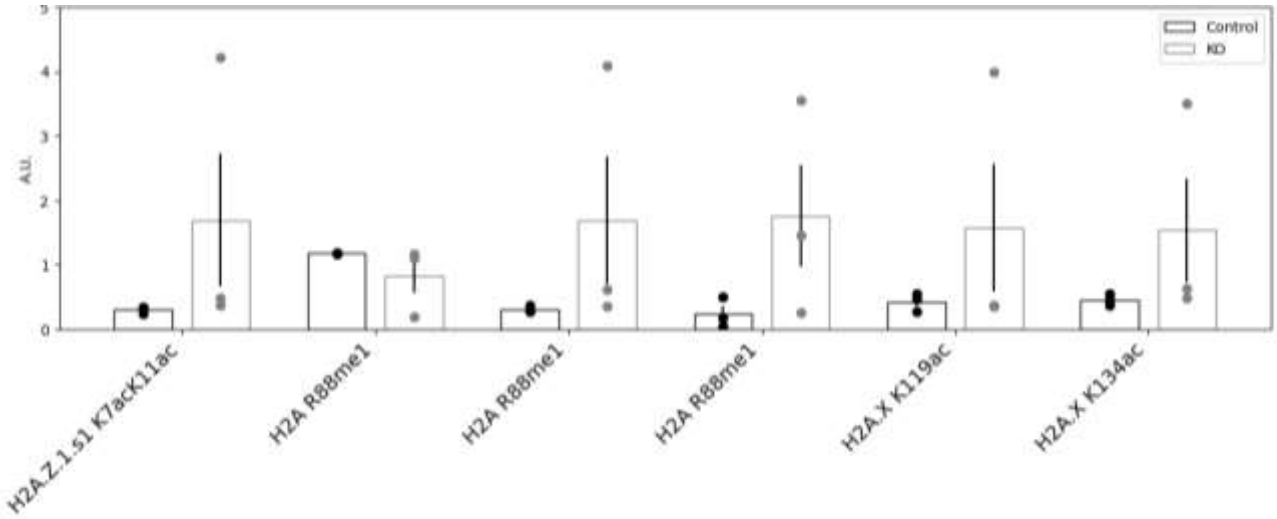

**H2B PTMs in RS**

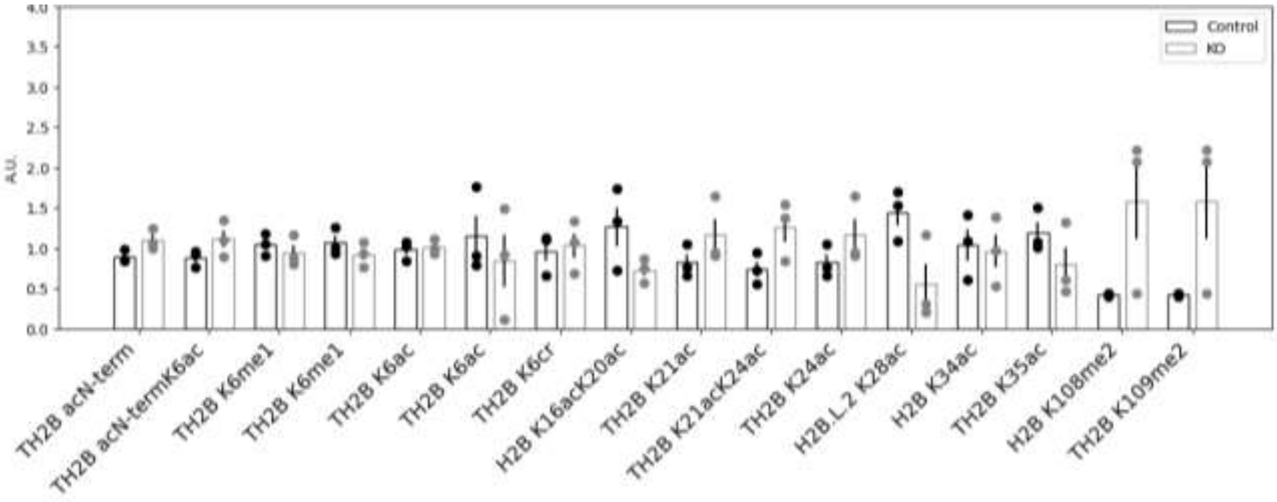

## S4F

**H3 PTMs in SPZ**

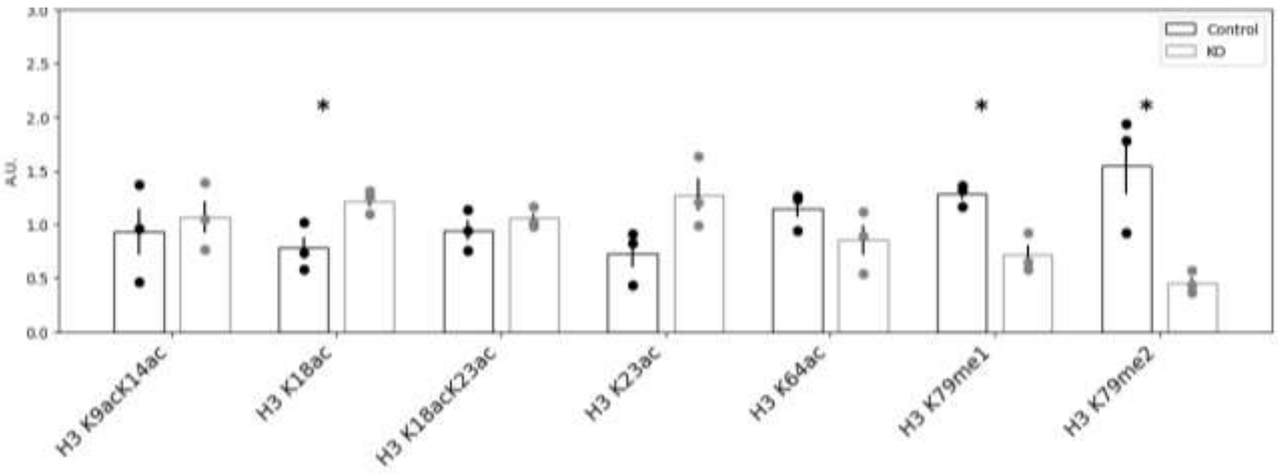

### Appendix Figure S5

## S5A

**Deregulated genes identified with DEseq2, with a Fold change > 1.5 and FDR < 0.05**

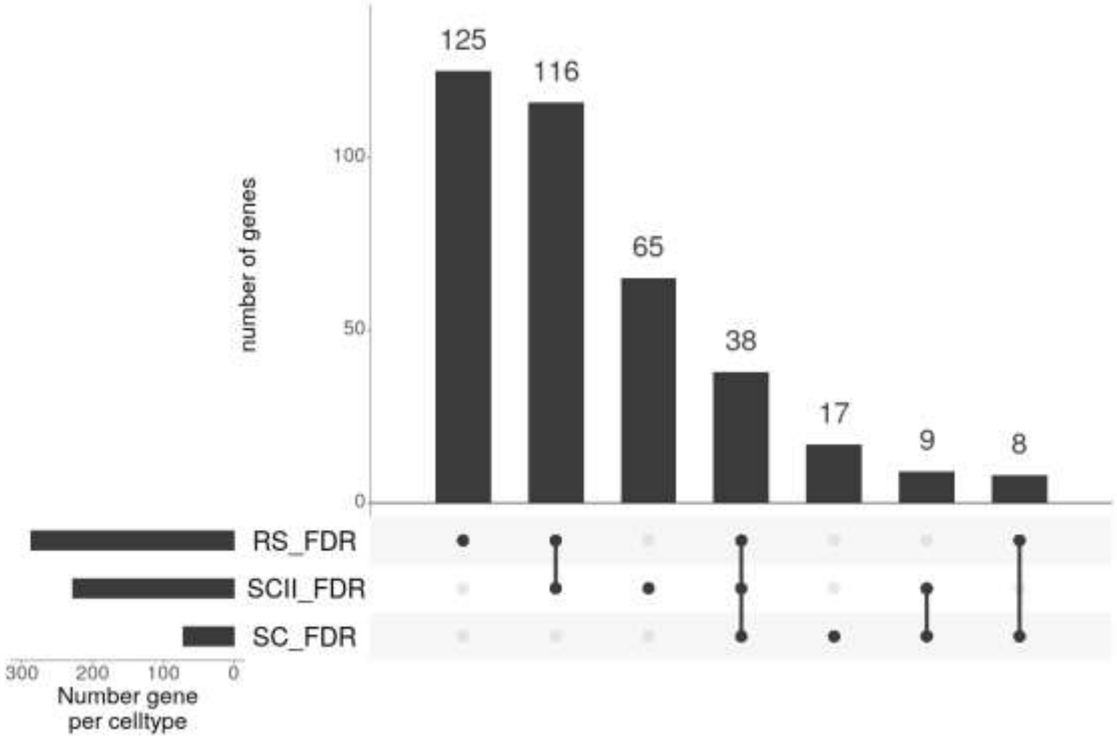

**Deregulated genes identified with DEseq2, with a Fold change > 1.5 and p.value < 0.05**

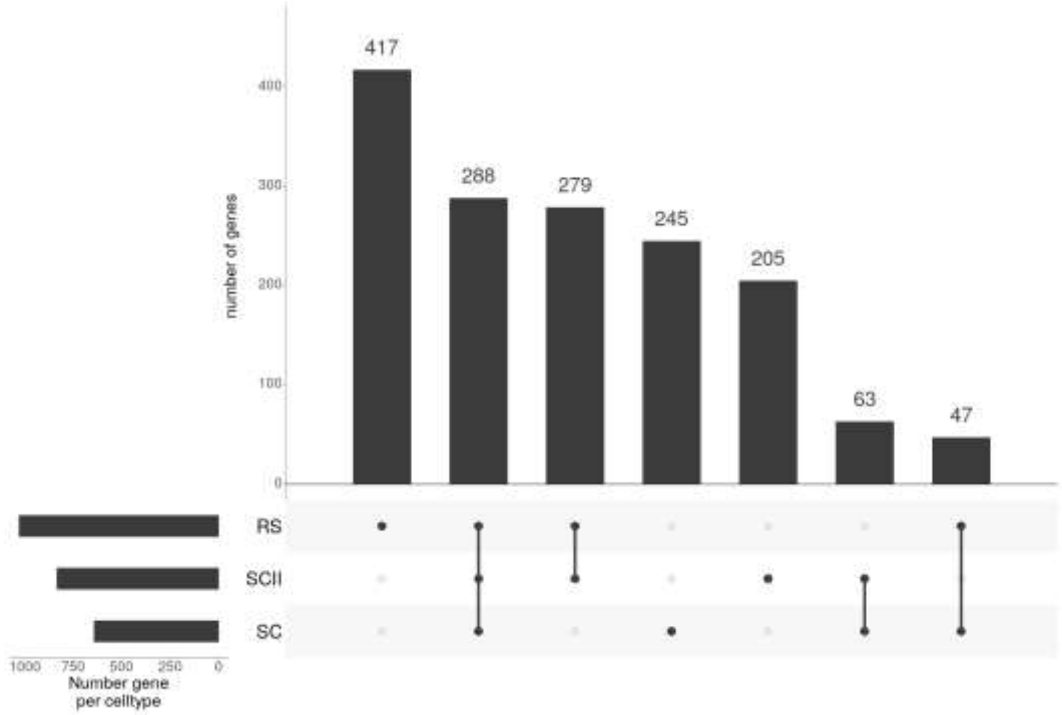

## S5B

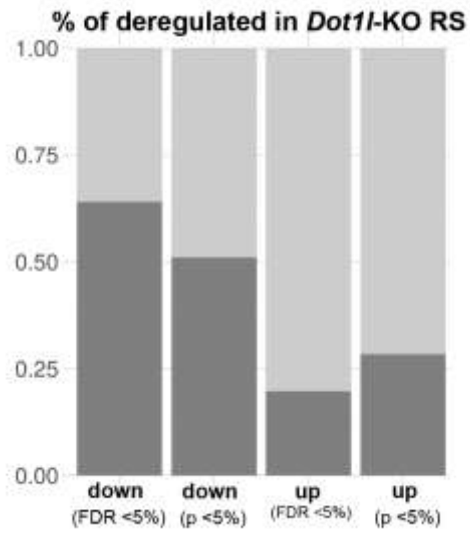

## S5C

| gene_name | logFC | logCPM | PValue | FDR | gl | H3K79me2 enrichment |
|---|---|---|---|---|---|---|
| *Slc10a3* | -1.26745740995669 | 0.999004290396908 | 0.00390006777314618 | 0.1545955595244 | Down-regulated | yes |
| *Slc6a8* | -2.46727862572467 | 0.52781470462503 | 7.72654921275588e-07 | 0.000364372904035715 | Down-regulated | yes |
| *Slc7a2* | -1.0665290941501 | 2.27338480123705 | 0.0150834110914719 | 0.387940102031817 | Down-regulated | no |
| *Slc12a1* | 1.3955625700193 | 0.681122253169346 | 0.000169908367523562 | 0.0174799371397939 | Up-regulated | no |
| *Slc13a4* | 1.54000399067509 | 2.76366740898566 | 4.27870310972635e-05 | 0.00657135254678289 | Up-regulated | no |
| *Slc18a2* | 1.25758541410194 | -0.793930612861824 | 0.00237996137428981 | 0.110886263192696 | Up-regulated | no |
| *Slc1a4* | 1.10997163431429 | 0.860924966152978 | 0.00103762614100171 | 0.0601552770604185 | Up-regulated | no |
| *Slc1a5* | 1.06878712737505 | 2.2418481205103 | 0.00176357549579316 | 0.0883504245162868 | Up-regulated | yes |
| *Slc25a23* | 0.926874320430452 | 2.4842629146059 | 0.00744458216430107 | 0.240033828551142 | Up-regulated | yes |
| *Slc25a37* | 0.83494237600095 | 2.8363663126306 | 0.0494170665013017 | 0.856402093077086 | Up-regulated | no |
| *Slc25a48* | 0.844791121775937 | 0.601566750878029 | 0.035227161221668 | 0.691260215461133 | Up-regulated | yes |
| *Slc26a11* | 1.65452313717142 | 1.74006812886016 | 0.000661650590507997 | 0.044270891766396 | Up-regulated | yes |
| *Slc28a2* | 0.882471439124064 | 0.348580484195426 | 0.0475801028844169 | 0.835942882356513 | Up-regulated | no |
| *Slc29a3* | 1.00130602592956 | 3.99897275785481 | 0.024747552656584 | 0.549198182271674 | Up-regulated | yes |
| *Slc37a3* | 0.864250867428252 | 1.06283489334379 | 0.0157601580162183 | 0.398622878559951 | Up-regulated | no |
| *Slc38a2* | 1.19621193912572 | 2.2217849521669 | 0.000373937024375424 | 0.0301146206327321 | Up-regulated | yes |
| *Slc38a8* | 1.03918406075334 | -0.0156846590541735 | 0.0201914826533337 | 0.472852642452675 | Up-regulated | no |
| *Slc45a1* | 1.75519777185066 | -0.803109163412898 | 0.000635433125120995 | 0.0430016682924086 | Up-regulated | no |

## S5D

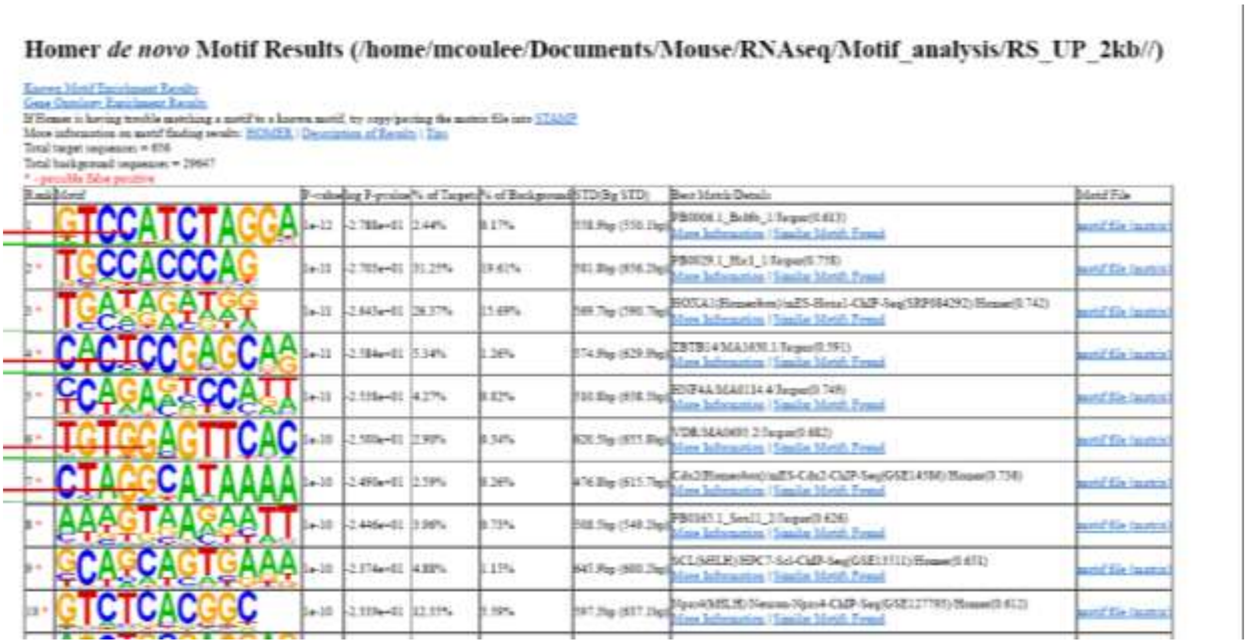

| gene_name | logFC | logCPM | PValue | FDR | gl | H3K79me2 enrichment |
|---|---|---|---|---|---|---|
| Cacna1e | 1.381540782 | 2.204690766 | 0.002726556 | 0.121318141 | Up-regulated | Yes |
| Gpsm1 | 1.115505708 | 1.018472463 | 0.002648857 | 0.119051423 | Up-regulated | No |
| Tmem74b | 1.021769193 | 0.247820597 | 0.013177832 | 0.354170753 | Up-regulated | No |
| Gata5 | 1.673646912 | -1.21958131 | 0.000102450 | 0.012941114 | Up-regulated | No |
| Padi1 | 1.006558376 | -0.52514070 | 0.027404915 | 0.589073299 | Up-regulated | No |
| Ptpn13 | 0.896515079 | 3.191552902 | 0.026370274 | 0.572363590 | Up-regulated | No |
| Megf8 | 0.875273628 | 5.337855272 | 0.023642554 | 0.532645802 | Up-regulated | No |
| Kctd21 | 0.837477696 | 0.230864728 | 0.024428532 | 0.543473782 | Up-regulated | No |
| Irf1 | 1.521852615 | 0.397338881 | 0.006670929 | 0.224722281 | Up-regulated | No |
| Lhfpl2 | 1.068600482 | -0.60887569 | 0.028270799 | 0.599286273 | Up-regulated | No |
| Grid1 | 1.387320345 | 2.179807865 | 0.000243895 | 0.022484676 | Up-regulated | Yes |
| Dscc1 | 1.087546432 | 3.096511586 | 0.000497737 | 0.036606338 | Up-regulated | No |
| Baiap2l2 | 0.874498322 | 0.165478650 | 0.034369221 | 0.684231999 | Up-regulated | No |
| Zfp52 | 1.351603089 | -0.97586843 | 0.001088020 | 0.062633064 | Up-regulated | No |
| Plin3 | 0.983536252 | 2.207539245 | 0.007122447 | 0.233024479 | Up-regulated | Yes |

### Appendix Table S1

| **Table S1** | | |
|---|---|---|
| **genotyping primers** | **sequence** | **PCR condition** |
| **Dot1l-F** | **GCAAGCCTACAGCCTTCATC** | **Annealing temperature = 58°C** |
| **Dot1l-R1** | **CACCGGATAGTCTCAATAATCTCA** | |
| **Dot1l-R2** | **GAACCACAGGATGCTTCAG** | |
| **STRA8CRE_oIMR9266F** | **AGATGCCAGGACATCAGGAACCTG** | **Annealing temperature = 62°C** |
| **STRA8CRE_oIMR9267R** | **ATCAGCCACACCAGACACAGAGATC** | |
| **Ymtxfp10_ internal control** | **TCACACAGATAAGAGGGTATTG** | |
| **Ymtxrp13_ internal control** | **GTTTTCCTATCAGGCCATCCA** | |

### Appendix Table S2

| **Table S2** | | | |
|---|---|---|---|
| **Western Blot - Primary antibodies** | | | |
| **Target** | **Manufacturer** | **Reference** | **Usage** |
| **DOT1L** | **CST** | **D4O2T** | **1/500 - 1/1000** |
| **Tubulin** | **Upstate** | **05-661** | **1/3000** |
| **H3K79me2** | **Abcam** | **ab3594** | **1/500** |
| **TH2B** | **Upstate** | **07-680** | **1/5000** |
| **H3** | **Abcam** | **ab 1791** | **1/3000 - 1/10000** |
| **H4K16ac** | **Upstate** | **07-329** | **1/750** |
| **Poly ac-H4** | **Millipore** | **06-866** | **1/250** |
| **TNP2** | **Santa Cruz Biotechnology** | **sc21106** | **1/250** |
| **PRM2** | **Briar Patch Biosciences** | **MAb-Hup2B** | **1/500** |
| **Western Blot - Primary antibodies** | | | |
| **Target** | **Manufacturer** | **Reference** | **Usage** |
| **Donkey anti-goat HRP** | **Santa Cruz Biotechnology** | **sc2020** | **1/5000** |
| **Goat anti-mouse HRP** | **ThermoFisher** | **31430** | **1/5000** |
| **Goat anti-rabbit HRP** | **ThermoFisher** | **31460** | **1/5000** |
| **Immunofluorescence-Immunohistochemistry** | | | |
| **Target** | **Brand** | **Reference** | **Usage** |
| **DOT1L** | **CST** | **D402T** | **1/100** |
| **DOT1L** | **Abcam** | **64077** | **1/100** |
| **H3K79me2** | **Diagenode** | **C15410051** | **1/100** |
| **GFP** | **ThermoFisher** | **CAB4211** | **1/100** |
| **Poly ac-H4** | **Millipore** | **06-866** | **1/200** |
| **TNP2** | **Santa Cruz Biotechnology** | **sc21106** | **1/50** |
| **Chromatin Immunoprecipitation** | | | |
| **Target** | **Brand** | **Reference** | |
| **H3K79me2** | **Diagenode** | **C15410051** | |
